# Inclusion Complexation of S-Nitrosoglutathione for Sustained NO Release and Reduced Device Infection

**DOI:** 10.1101/2022.10.03.510719

**Authors:** Wuwei Li, Danyang Wang, Ka Un Lao, Xuewei Wang

## Abstract

S-nitrosoglutathione (GSNO) is a non-toxic nitric oxide (NO)-donating compound that occurs naturally in the human body. The use of GSNO to deliver exogenous NO for therapeutic and protective applications is limited by the high lability of dissolved GSNO in aqueous formulations. In this paper, we report a host-guest chemistry-based strategy to modulate the GSNO reactivity and the NO release kinetics. Cyclodextrins (CDs) are host molecules that are typically used to encapsulate hydrophobic guest molecules into their hydrophobic cavities. However, we found that CDs form inclusion complexes with GSNO, an extremely hydrophilic molecule with a solubility of over 1 M at physiological pH. More interestingly, the host-guest complexation reduces the decomposition reactivity of GSNO in the order of αCD > γCD > hydroxypropyl βCD. The lifetime of 0.1 M GSNO is increased to up to 15 days in the presence of CDs at 37°C, which is more than twice the lifetime of free GSNO. Quantum chemistry calculations indicate that GSNO in αCD undergoes a conformational change that significantly reduces the S-NO bond distance and increases its stability. The calculated S-NO bond dissociation enthalpies of free and complexed GSNO well agree with the experimentally observed GSNO decomposition kinetics. The NO release from GSNO-CD solutions, compared to GSNO solutions, has suppressed initial bursts and extended durations, enhancing the safety and efficacy of NO-based therapies and device protections. In an example application as an anti-infective lock solution for intravascular catheters, the GSNO-αCD solution exhibits potent antibacterial activities for both planktonic and biofilm bacteria, both intraluminal and extraluminal environments, both prevention and treatment of infections, and against multiple bacterial strains including a multidrug-resistant strain. In addition to solutions, the inclusion complexation also enables the preparation of GSNO hydrogels with enhanced stability and improved antibacterial efficacy. Since methods to suppress and control the GSNO decomposition rate are rare, this supramolecular strategy provides new opportunities for the formulation and application of this natural NO donor.

## 1. Introduction

Nitric oxide is a multifunctional gaseous radical that plays pivotal roles in a wide range of physiological and pathophysiological processes, such as vasodilation, coagulation, inflammation, neurotransmission, host defense, and wound healing [1–5]. Inspired by the biological functions of endogenous NO, exogenous NO has been employed as a therapeutic agent for pulmonary, cardiovascular, neurological, and renal diseases associated with NO deficiencies [6,7]. NO has also been released or generated from medical implants and dressings to protect against thrombosis, infection, and inflammation [8–10]. There are three main avenues of NO delivery. First, NO gas is inhaled into the respiratory system, which is an FDA-approved therapy for hypoxic respiratory failure associated with pulmonary hypertension [11]. Second, solid NO donors are embedded in nanoparticles and polymers, allowing for controlled release of NO from storable nanocarriers and devices [12]. A large number of publications focus on the functionalization of polymeric devices such as catheters, grafts, cannulas, and sensors with NO donors or NO-donating moieties to reduce device-associated complications [9,10,12–15]. Third, aqueous solutions, suspensions, and hydrogels containing NO donors are suited to be administrated via intranasal, intramuscular, intracameral routes, or topically for therapeutic purposes [16–19]. They have also been employed as filling solutions for implants such as catheters [20–25], representing a unique method to release protective NO without modifying the polymers of medical implants.

Compared to NO delivery from NO tanks/generators and embedded solid NO donors, controlled release of NO from aqueous formulations is more challenging due to the low solubility, high reactivity, and toxicity of many NO donors. For example, sodium nitroprusside and N-diazeniumdiolates pose a risk of toxicity due to their degradation byproducts, including thiocyanate and nitrosamines [26,27]. S-Nitroso-N-acetylpenicillamine, a commonly used synthetic S-nitrosothiol type NO donor, has a low aqueous solubility of only 2.1 mg/mL (< 0.01 M) in water [28]. GSNO is a natural NO carrier and transporter circulating in the blood and occurring within the cytoplasm of cells [29]. Compared to other donors, GSNO is especially suitable for water-based drug formulations. As the S-nitrosated derivative of tripeptide glutathione, GSNO has an aqueous solubility of 0.075 M at low pH and > 1 M at the physiological pH. *In vivo* studies did not reveal any toxicities of this natural NO donor when administrated in humans, dogs, and rats at appropriate doses [30,31]. There have been nearly 20 clinical trials using GSNO as a therapeutic drug, further confirming its safety [29,32]. However, one long-standing and well-recognized challenge of NO delivery via GSNO solutions is the high GSNO reactivity that leads to “rapid and often unpredictable” rates of NO generation in medical applications [32]. GSNO readily decomposes under heat (e.g., 37 °C) and light, and in the presence of catalysts/reactants such as metal ions, thiols, ascorbic acid, enzymes, and proteins [29,33]. Its decomposition rate is also concentration dependent via a thyil radical-based autocatalysis mechanism [34,35].

A high NO donor concentration is needed in many preventative and therapeutic applications to release adequate NO from a limited volume of solution. Simply dissolving GSNO in an aqueous solution may not work due to its high reactivity at high concentrations at the physiological temperature [35]. The total NO release duration is usually too short to provide sustainable benefit and the high initial burst release causes cytotoxicity and multiple adverse effects [15,36]. Indeed, the biological function of NO is often bidirectional and highly dependent on its concentration [37]. Consequently, the modulation of the GSNO reactivity for sustained release of NO with reduced initial burst is a key to the successful implementation of GSNO solutions in many medical applications. The encapsulation of GSNO in suspended nanocarriers based on poly(methyl)methacrylate, alginate, chitosan, and liposomes has been reported with the aim of controlling the GSNO decomposition [12,19,38,39]. However, these nanocarriers usually have low drug loadings and the actual NO release duration is only several hours to several days.

The formation of host-guest inclusion complexes is a common strategy in drug formulation to improve the physicochemical stability, solubility, dissolution rate, and bioavailability of drugs [40]. CDs are one of the most widely used families of host molecules for drug inclusion due to their low toxicity, wide availability, and low cost. Naturally occurring α, β, and γCDs consist of 6, 7, and 8 glucopyranose units, respectively, and differ in their cavity size and solubility [41]. These parent CDs can be further chemically modified to various derivatives to possess more diversified physicochemical and biopharmaceutical properties as well as drug encapsulation capabilities [42]. CDs have a cone-shaped structure with a hydrophilic exterior and a lipophilic cavity and are, therefore, typically employed to encapsulate hydrophobic drugs via intermolecular forces [41]. Indeed, nearly all commercial drugs complexed with CDs, such as prostaglandin, dexamethasone, piroxicam, β-oestradiol, itraconazole, voriconazole, and ziprasidone are insoluble or sparingly soluble in water (solubility of several mg/mL or below) [43]. β-CD forms an inclusion complex with nitroglycerin, an NO-donating organic nitrate with an aqueous solubility of only 1.4 mg/mL, to enhance the drug stability in preventing angina pectoris (“Nitropen”) [44,45]. Hydroxypropyl-βCD was found to stabilize S-nitroso-N-acetylpenicillamine at pH ∼3 under which the aqueous solubility is only 2.1 mg/mL [46].

We are interested in regulating the decomposition of concentrated GSNO at physiological pH via host-guest interactions. Given that CDs are mostly suited to include hydrophobic drugs in their apolar cavities, it is not surprising that the inclusion complexation of highly hydrophilic GSNO with CDs has not been reported, although both molecules have been intensively studied in pharmaceutical sciences. Herein, we demonstrated that GSNO at physiological pH forms inclusion complexes with appropriate CDs such as αCD. The rate of thermal decomposition, photodecomposition, and reactions with biological molecules of encapsulated GSNO is significantly reduced compared to free GSNO. NO release in a more sustained and steady fashion has been obtained. In an example application, the GSNO-CD solutions used as lock solutions of intravascular catheters are highly effective in preventing and killing planktonic and biofilm bacteria. Notably, this work differs from previous reports on the modification of CDs with NO-donating moieties via covalent bonding, which neither used CDs as host molecules nor focused on regulating the NO release kinetics [47,48].

## 2. Experimental methods

### 2.1 Chemicals and reagents

Sodium phosphate dibasic (Na_2_HPO_4_), sodium hydroxide (NaOH), ethylenediaminetetraacetic acid disodium salt dihydrate (EDTA·2Na), L-glutathione reduced (GSH), cysteine, L-ascorbic acid, bovine serum albumin (BSA), and sodium nitrite were purchased from MilliporeSigma. Luria-Bertani (LB) broth powder, agar, αCD, Dulbecco’s modified Eagle’s medium (DMEM), fetal bovine serum (FBS), and penicillin/streptomycin were purchased from Thermo Fisher Scientific. Gamma CD and 2-hydroxypropyl βCD (HP βCD) were purchased from Tokyo Chemical Industry and Cayman Chemical Company, respectively. Other CD derivatives were purchased from Cyclolab Ltd. Bacterial strains including *S. aureus* (25923), methicillin-resistant *S. aureus* (MRSA, BAA-2312), *S. epidermidis* (12228), *E. coli* (53496), and *P. aeruginosa* (baa-744) as well as murine fibroblast cell line (L929) were purchased from the American Type Culture Collection (ATCC). The CellTiter 96® AQueous One Solution Cell Proliferation Assay (MTS) kit was obtained from Promega Corporation.

### 2.2 GSNO synthesis and preparation of NO release solutions

GSNO was synthesized by nitrosation of reduced L-glutathione in acidified nitrite as previously reported [49]. After collecting GSNO precipitates and washing them using cold water and acetone, GSNO powder was vacuum dried for 4 h and stored at -20°C. GSNO powders were dissolved in Na_2_HPO_4_ to prepare GSNO solutions. One hundred µM EDTA was added to each solution as a chelating agent to mask Cu^2+^ before the solution pH was finally adjusted to 7.4 via NaOH. For solutions with reactive agents, BSA, ascorbic acid, or cysteine was added before the pH adjustment. To prepare GSNO-CD solutions, appropriate amounts of CD powders were dissolved into the GSNO solutions prior to the pH adjustment.

### 2.3 Measurement of GSNO decomposition via UV–vis absorption spectroscopy

Aqueous GSNO solutions were contained in 1.5 mL disposable polystyrene cuvettes. The cuvette was caped and further sealed with parafilm. The concentration of GSNO during storage at 37 °C in the dark was monitored by a UV-vis spectrophotometer (Go Direct® Fluorescence/UV-VIS Spectrophotometer) at a wavelength of 545 nm. A 4 W LED white light was used as the light source for the photodecomposition experiment at room temperature. The distance between the light source and the cuvettes was approximately 10 cm. All GSNO decomposition experiments were triplicated, and the GSNO concentration was expressed as mean ± standard deviation.

### 2.4 Measurement of NO release

The NO release was quantified by an ECO PHYSICS NO analyzer (nCLD 66). Two μL of castor oil was added to NO release solutions as an antifoam agent when 3 mL GSNO or GSNO-CD solutions were tested in an amber glass cell at 37 °C. Humidified air was vacuumed into the solution to carry NO continuously into the chemiluminescence detection chamber at a flow rate of 100 cm^3^/min. The solutions were stored in sealed amber glass vials between measurements at 37 °C. To monitor the NO release from catheters, solutions were filled into 2.5 cm-long medical-grade silicone tubes (HelixMark® 60-011-07, 1.58 mm ID, 2.41 mm OD) that were sealed on both ends with plastic rods. The filled tubes were placed into the amber glass cell containing 5 mL of PBS with 0.1 mM EDTA (PBSE) at 37 °C. N_2_ was used as the carrier gas at a flow rate of 100 cm^3^/min to introduce NO into the analyzer. Filled tubes were stored in 10 mL PBSE at 37 °C in the dark between measurements. The soaking PBSE solution was refreshed after each measurement.

### 2.5 Preparation of agar hydrogel containing GSNO

Two grams of agar were dissolved in 100 mL of 0.1 M phosphate buffer at pH 7.4 at 80°C for 3 h with stirring. GSNO solutions at 0.2 M with or without 0.2 M αCD in 0.1 M phosphate buffer at pH 7.4 were prepared separately. Then, 0.5 mL GSNO solution and 0.5 mL dissolved agar were mixed and put into an ice bath to facilitate gelation.

### 2.6 Quantum chemistry calculations

The free GSNO molecule (both carboxylic groups deprotonated) and its binding with α, HP β, and γCD were optimized at the level of r^2^SCAN-3c [50] with the water solvent effects considered by using the conductor-like polarizable continuum model (CPCM) [51]. The composite electronic-structure method r^2^SCAN-3c has been shown to obtain reliable binding energies even at large noncovalent complexes [52] and should provide reliable binding energies for GSNO with CDs. The frequency calculations at room temperature have been employed to obtain thermal corrections and confirm all structures as minima. All calculations were performed using version 5.0.3 of the ORCA program [53].

### 2.7 *In vitro* antibacterial experiments

Both biofilm and planktonic bacteria were quantified to evaluate the antibacterial activities of the proposed method. LB medium (25 mg/mL) was used to culture all bacterial strains overnight before performing the antibacterial tests. Segments of medical-grade silicone tubes were all autoclaved. Lock solutions were freshly prepared using the sterilized buffer and filtered through a 0.22 μm PES syringe filter. The biofilms on the silicone surface were dip-rinsed 5 times in PBS and detached by 1-min vertexing, 1-min sonication (20% power of a 150W probe sonifier, Branson Ultrasonics), and another 1-min vertexing for plate counting. When quantifying the planktonic bacteria, the suspension was homogenized by 1-min vertexing. Bacterial suspensions were serially diluted using sterile PBS prior to inoculating the 1.5 wt% agar LB plates. After plate incubation at 37°C overnight, colony-forming units (CFU) of bacteria were counted and converted to CFU/cm^2^ for biofilm quantification and CFU/mL for planktonic bacteria quantification. All bacterial tests were triplicated. Data were reported as mean ± standard deviation. The statistical significance between groups was determined using a student’s t-test.

#### 2.7.1 Extraluminal bacteria assessment

The overnight *S. aureus* culture was diluted by LB (0.25 mg/mL) medium to approximately 10^3^ CFU/mL. Silicone rubber tubing was cut into 2.7 cm segments and one end was sealed with RTV silicone rubber glue. The silicone segment was filled with a lock solution and then transferred to a 15-mL centrifuge tube with 1.8 mL bacterial culture. The top end of the catheter segment was left open and above the liquid culture. Centrifuge tubes were capped loosely and incubated statically at 37°C in an incubator. The catheter segments were dip-rinsed 5 times in sterile PBS to remove the loosely attached bacteria before the broth was refreshed once daily. At the end of the experiment, the lock solution was completely extracted and discarded. The empty silicone tube was thoroughly rinsed and transferred to 5 mL PBS to perform the biofilm evaluation experiment detailed above. Glutathione (0.25 mg/mL) is present in the LB medium to mimic the thiol environment of blood. To simulate the lock solution therapy for bacteria eradication, silicone segments were capped with plastic plugs at both ends. Bacterial biofilms were formed on the exterior surface after incubating the sealed segment in *S. aureus* culture for 48 h at 37°C. Then, the catheter lumen was filled with different lock solutions. The infected silicone tubes were transferred to 15-mL centrifuge tubes with PBS and incubated for 24 or 48 h at 37 °C. The viable bacteria on the outer surface were counted following the treatment.

#### 2.7.2 Intraluminal bacteria assessment

αCD, GSNO, and GSNO+αCD solutions were prepared using 0.1 M Na_2_HPO_4_ buffered LB broth and adjusted to pH 7.4. Then, the overnight bacterial cultures were 100-fold diluted by the buffered LB broth, αCD, GSNO, or GSNO+αCD solutions. Silicone tubing was cut into 2 cm segments and then cut longitudinally into two halve to fully expose all surfaces, mimicking the interaction between the inner silicone surface and the lock solution. The silicone pieces were soaked into the prepared solutions with bacteria and statically incubated for 3 days at 37 °C. Viable biofilm and planktonic cells were evaluated by plate counting. To simulate the lock therapy for intraluminal bacterial biofilm, sterile silicone pieces were exposed to *S. aureus* for 48 h at 37°C to grow biofilm. Then, these pieces were treated with different lock solutions for another 24 h followed by the biofilm evaluation process described above.

#### 2.7.3 Agar diffusion assay for the hydrogel containing GSNO

The overnight *S. aureus* culture was diluted to a density of 0.1 OD_600_ in PBS. Then, the diluted bacterial inoculum was uniformly spread on an LB-agar plate using a sterile cotton swab. The plate was marked as four separate portions. Forty μL of warm liquid agar with only buffer, GNSO, αCD, or GSNO-αCD (see section 2.5) was added onto a portion of the inoculated plate twice. The agar solution was gelated almost instantly due to the temperature decrease to form a small circular hydrogel area. The bacterial growth was visually examined after 24-h incubation at 37°C.

### 2.8 Cytotoxicity test

L929 murine fibroblast cells were suspended in complete media (DMEM with 1% penicillin/streptomycin and 10% FBS) before being seeded in a 24-well plate at a concentration of 5×10^5^ cells per well. After 24 h of incubation, the attached cells in each well were incubated in fresh media containing a sterilized 1-cm silicone tube (HelixMark® 60-011-07) filled with a lock solution with buffer, GSNO, or GSNO-αCD and with both ends sealed. MTS assays were performed following the manufacturer’s procedure after an additional 24-h incubation. Absorbance at 490 nm was measured by a Cytation 3 imaging plate reader (Biotek).

## 3. Results and discussion

### 3.1 Effect of CDs on the thermal decomposition of GSNO in aqueous solutions and hydrogels

GSNO decomposes to liberate NO at the physiological temperature via thermal cleavage of its S–NO bond. Using an optimized buffer system according to our previous work [54], 0.1 M GSNO lasts 6 days in 0.1 M PBSE at pH 7.4 and 37°C (Figure 1A, black line). However, 66% of GSNO degrades within the first 24 h, suggesting an initial burst release of NO. When the GSNO solution contains 0.1 M CD, the GSNO stability is enhanced in the order of αCD > γCD > HP βCD (hydroxypropyl βCD). The solubility of parent βCD in water is only 16.5 mM, which is inadequate to complex a high concentration of GSNO. So, we chose a commonly used derivative, HP βCD as a more soluble substitute. An equal mole of αCD increases the GSNO lifetime to 11 days, which is nearly twice the lifetime of GSNO only. The GSNO decay in the first 24 h also decreases to only 39%, suggesting a much lower initial decomposition reactivity in the presence of the hosting αCD. As shown in Figure 1B, the stabilization effect is dependent on the molar ratio of the host and guest molecules. When the αCD concentration is close to its maximum solubility (0.15 M), the GSNO decomposition rate is further reduced. To overcome the solubility limitation, multiple CDs may be used at high concentrations together to inhibit the GSNO decomposition. The GSNO lifetime is increased to 15 days with a first-day decay of only 22% in the presence of 0.15 M αCD and 0.1 M γCD (Figure 1B, purple line). The lifetime and first-day decay are improved by 2.5 and 3 times, respectively. When the concentration of GSNO is reduced to 0.05 M, similar trends of stabilization were observed in the presence of different CDs and different concentrations of αCD (Figure 1C). The GSNO lifetime is 17 days when 0.05 M GSNO and 0.15 M αCD are dissolved in 0.05 M PBSE at pH 7.4 at 37°C, as opposed to only 7 days without CDs. The stabilization effect is also pronounced at room temperature. As shown in Figure 1D, the half-life of GSNO is ∼ 20 days in the presence of 0.15 M αCD, which is 4 times that of only GSNO.

**Figure 1.**
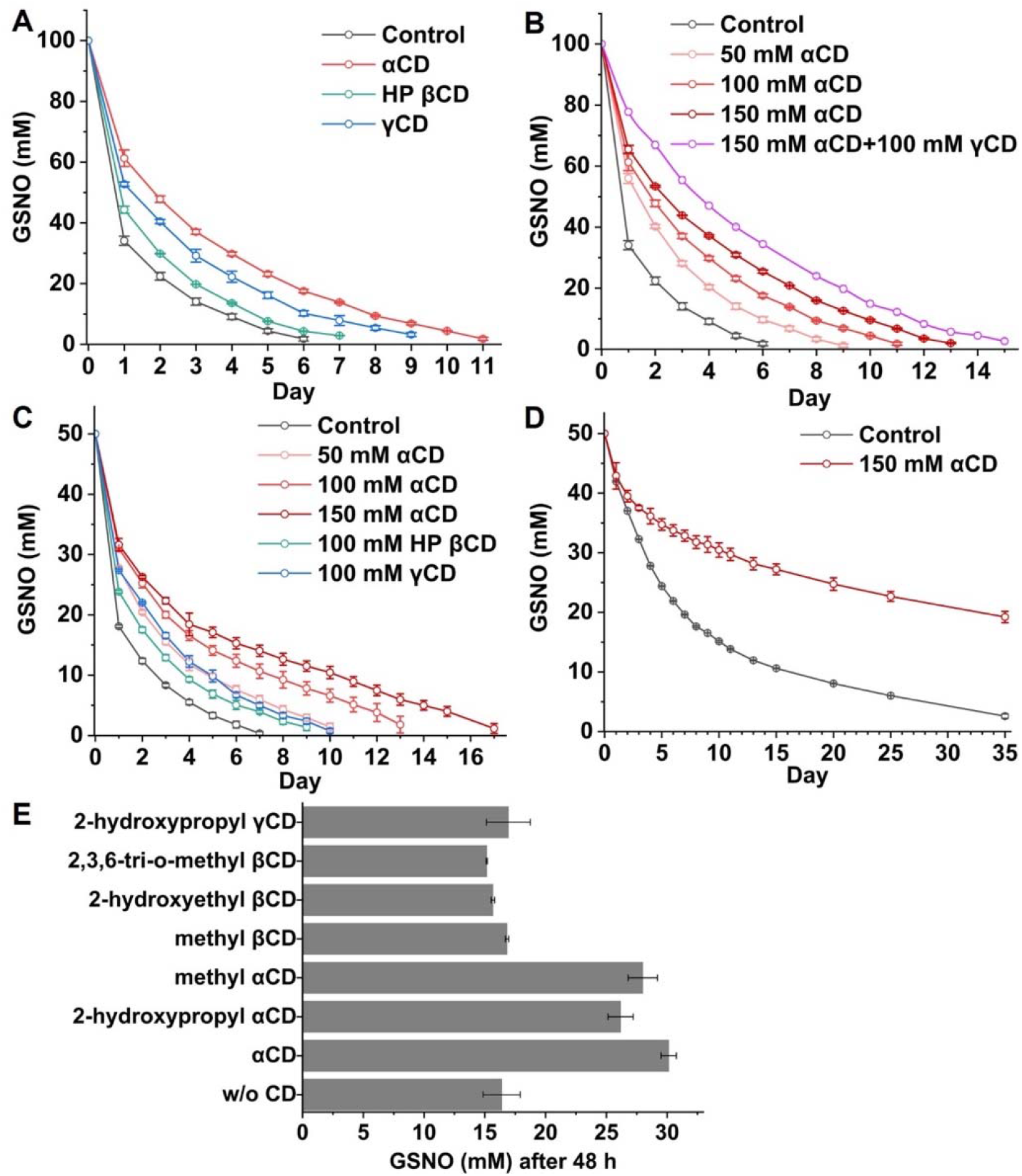
(A) Decay curves of 0.1 M GSNO in 0.1 M PBSE at pH 7.4 and 37°C without CD and with 0.1 M αCD, HP βCD, and γCD. (B) Decay curves of 0.1M GSNO in 0.1 M PBSE at pH 7.4 and 37°C with 0, 0.05, 0.1, 0.15 M αCD and 0.15 M αCD+0.1 M γCD. (C) Decay curves of 0.05 M GSNO in 0.05 M PBSE at pH 7.4 and 37°C with and without CDs. (D) Decay curves of 0.05 M GSNO in 0.05 M PBSE at pH 7.4 at room temperature with or without αCD. (E) The remaining GSNO after 48-h 37°C storage of 0.05 M GSNO solutions in the presence of 0.1 M various CD derivatives.

We further examined six other CD derivatives in a 2-day experiment and found that parent αCD is still the best (Figure 1E). If we group the derivatives based on their parent CDs, the stabilization effect also follows the order of αCD > γCD > βCD. Generally, βCD and its derivatives are the most commonly used and successful complexing agents in pharmaceutical formulations as their cavities match the size of many drugs [43]. Our results indicate that αCD is most effective in modulating the GSNO reactivity. αCD derivatives, including methyl αCD and 2-hydroxypropyl αCD, also show a significant stabilization effect, but these derivatives are more expensive than the parent αCD and therefore not further studied in this work.

The decomposition of each GSNO molecule liberates one NO. Parent CDs, alkylated CDs, and hydroxyalkylated CDs examined in this study are not expected to react with NO and consume NO. Therefore, the decay kinetics of GSNO based on absorption spectroscopy should agree with the NO generation kinetics. To confirm this, NO release from 3 mL of 0.1 M GSNO solution (3×10^−4^ moles GSNO) with and without αCD is tested using a chemiluminescence-based NO analyzer. As shown in Figure 2, GSNO and GSNO-αCD solutions release NO for 6 and 11 days, respectively, which is completely consistent with the absorbance-based GSNO decay data. The cumulative NO release is calculated to be 2.48×10^−4^ and 2.36×10^−4^ moles, respectively, confirming that the presence of αCD does not consume NO.

**Figure 2.**
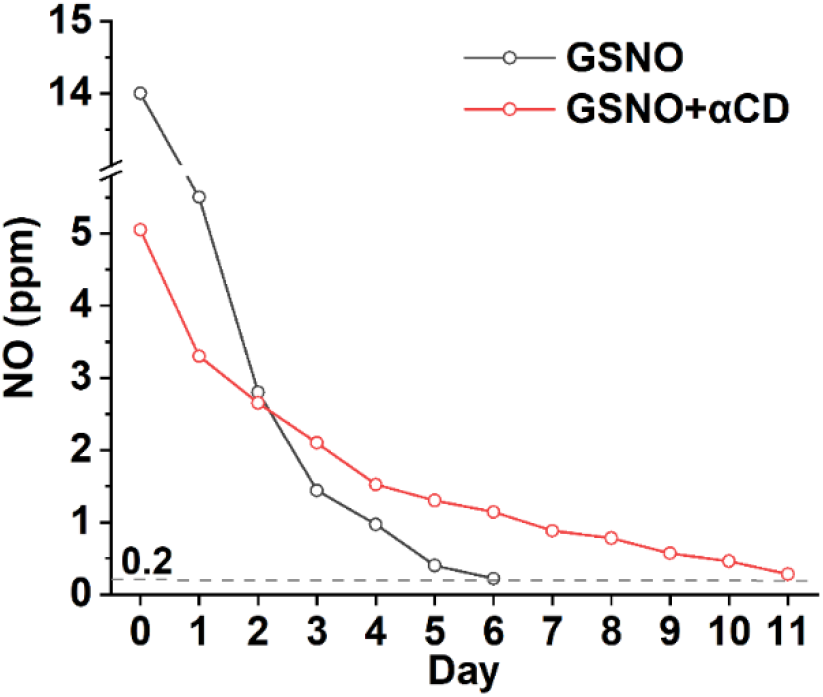
NO release from 3 mL of 0.1 M GSNO solutions with or without an equal mole of αCD at 37°C.

NO-releasing hydrogels have been shown to promote wound healing in topical applications [55], enhance tumor immunotherapy as injectables [56], and protect medical implants as coatings [8]. Therefore, we studied the effect of αCD on the stability of GSNO in hydrogel using agar hydrogel as an example. As shown in Figure 3, the reddish color of GSNO in 1% agar hydrogel fades in less than 2 days at 37°C, whereas the color lasts more than 6 days in the presence of an equal mole of αCD, suggesting that CD is a versatile host molecule to enhance GSNO stability in various water-based media. GSNO solutions have also been used in the preparation of NO-releasing water-in-oil emulsions [19] and liposomes [39], where the addition of CDs is also expected to modulate the GSNO decomposition and NO release.

**Figure 3.**
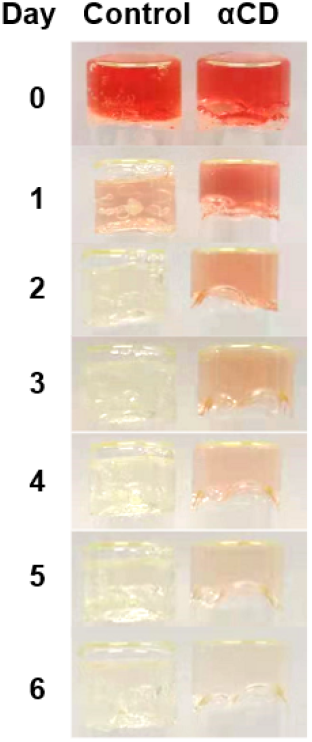
Photos of agar hydrogels containing 0.1 M GSNO with and without an equal mole of αCD after different days of storage at 37°C.

### 3.2 Computational investigation of GSNO-CD complexes

To understand the host-guest interaction between αCD, HP βCD, and γCD with GSNO, r^2^SCAN-3c has been used to calculate their structures, binding free energies, S-NO bond dissociation enthalpies, and the corresponding S-NO bond distances. In order to obtain the most stable structure for each binding complex, a wide range of candidate structures have been considered. The final binding free energies for GSNO with αCD, HP βCD, and γCD are -0.9, -8.2, and -9.0 kcal/mol, respectively, with the corresponding geometries shown in Figure 4. GSNO binds stronger with γCD due to the suitable size of γCD to hold GSNO where the interactions come from dispersion interactions of GSNO with the inner bowl of γCD and three hydrogen bonds of GSNO with the bowl edge. The binding is slightly weaker for GSNO with HP βCD which has a smaller size because a part of GSNO starts to leave the bowl and gives up the binding with the inner bowl. The binding is largely reduced for GSNO with the smallest α-CD since its bowl diameter is too small to fit GSNO. To compensate for the lack of binding with the inner bowl in α-CD, GSNO changes its conformation to form four hydrogen bonds with the bowl edge to maximize its binding. This can be seen by comparing the energy of GSNO in CD with the free GSNO molecule. GSNO conformations in αCD, HP βCD, and γCD is 10.4, 2.4, and 0.9 kcal/mol, respectively, higher in energy as compared with the free GSNO molecule. However, the S-NO bond distance is the shortest in the highest energy GSNO conformation in α-CD. The S-NO bond distance of GSNO in αCD, HP βCD, and γCD is 1.823, 1.852, and 1.879 Å, respectively. The S-NO distance of GSNO in α-CD is significantly smaller than that in the free GSNO molecule (1.875 Å), which indicates the stronger S-NO bond of GSNO in α-CD to defer the NO release. The corresponding S-NO bond dissociation energy (BDE) of GSNO in αCD, HP βCD, and γCD is 34.1, 31.0, and 33.0 kcal/mol, respectively. As compared with the S-NO BDE of the free GSNO molecule (31.3 kcal/mol), GSNO binding with αCD enhances the S-NO bond most and binding with γCD also enhances the S-NO bond. The S-NO BDE in the order of αCD > γCD > HP βCD agrees perfectly with the experimentally observed GSNO decomposition rates. In short, the decomposition reactivity of CD-complexed GSNO is determined by the BDE of S-NO after the inclusion of complexation and conformational change and cannot be simply explained by binding free energy trend, γCD > HP βCD > αCD.

**Figure 4.**
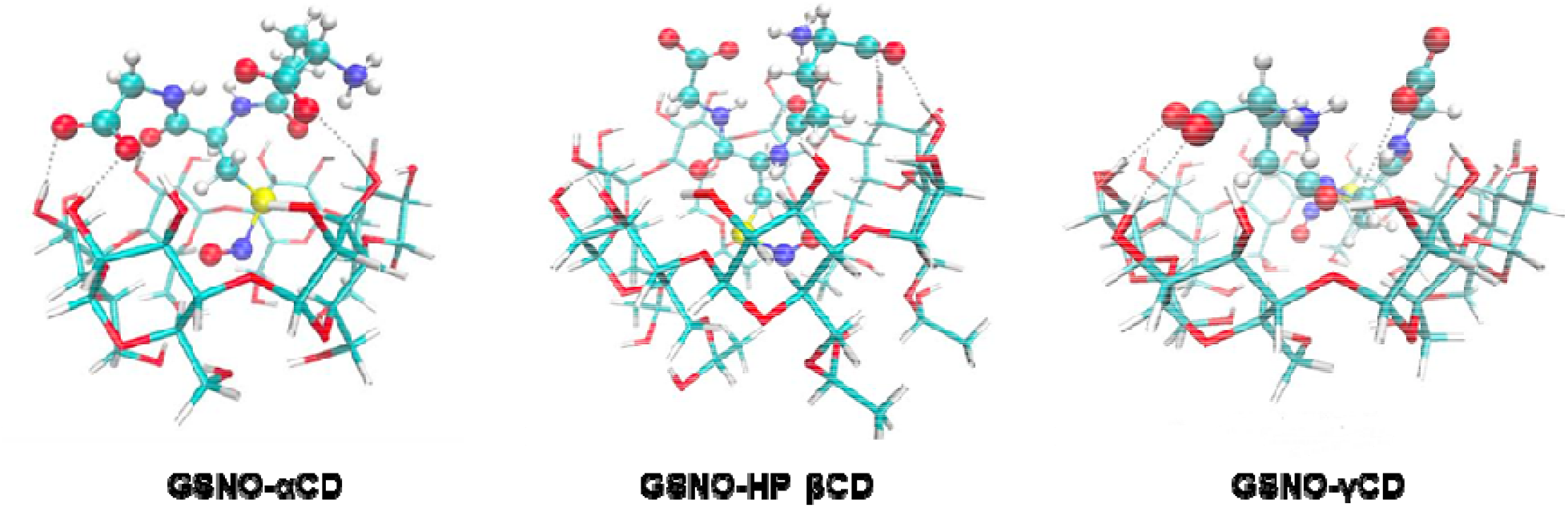
Calculated GSNO-CD complexes at the physiological pH. The ball-and-stick models represent negatively charged GSNO molecules with both carboxylic acids dissociated. The wire models represent different CDs. Gray dotted lines indicate the hydrogen bonds between GSNO and CDs. The coloring system is as follows: white (hydrogen), cyan (carbon), dark blue (nitrogen), red (oxygen), and yellow (sulfur).

### 3.3 Stabilization effect of αCD on GSNO in the presence of light and biologically relevant reactants

When GSNO solutions are mixed or in contact with biological fluids, the GSNO decomposition and NO release are expected to be faster due to the presence of enzymes such as GSNO reductase and γ-glutamyl transpeptidase, reactants such as ascorbic acid, thiols, metal ions, and proteins, as well as NO scavengers like hemoglobin [29,33]. Therefore, we further studied the effect of CD complexation on these reactions. Cysteine and ascorbate cause reductive cleavage of the S–N bond in GSNO and other S-nitrosothiols. As shown in Figure 5A and 5B, the addition of 10 mM cysteine and L-ascorbic acid reduces the lifetime of 0.1 M GSNO to 2-3 days in 0.1 M PBSE at pH 7.4, much shorter than the 6-day lifetime without these reductive compounds. However, the addition of 0.1 M αCD remarkably reduces the decomposition rate and approximately doubles the GSNO lifetime. Although the actual cysteine and ascorbate concentration *in vivo* is lower than the used 10 mM, multiple reductive reactants coexist in real biofluids such as blood and may accelerate the GSNO decomposition together. The purpose of this experiment is to clearly show that the host-guest interaction modulates the reaction of GSNO in the presence of biologically relevant reactants instead of implying specific reaction rates in real biological environments. S-transnitrosylation of proteins, including albumin, by GSNO is one of the *in vivo* generation mechanisms of S-nitrosated proteins, such as S-nitrosoalbumin, the most abundant circulating NO carrier in the blood [57]. To explore whether CDs could regulate the reaction of GSNO with proteins, BSA was added to GSNO solutions with and without αCD. As can be seen from Figure 5C, 41% GSNO is left after 24 h of storage at 37 °C in the presence of 0.1 M αCD, whereas only 21% is left in the control group. Photolytic decomposition of GSNO and other S-nitrosothiols is another widely studied reaction for NO generation. The presence of αCD leads to a nearly 2-fold decrease in the decomposition rate of GSNO in a 4-h experiment under LED white light (Figure S1). Collectively, the inclusion complexation of GSNO by CD is an effective method to reduce GSNO reactivity in various reactions and conditions.

**Figure 5.**
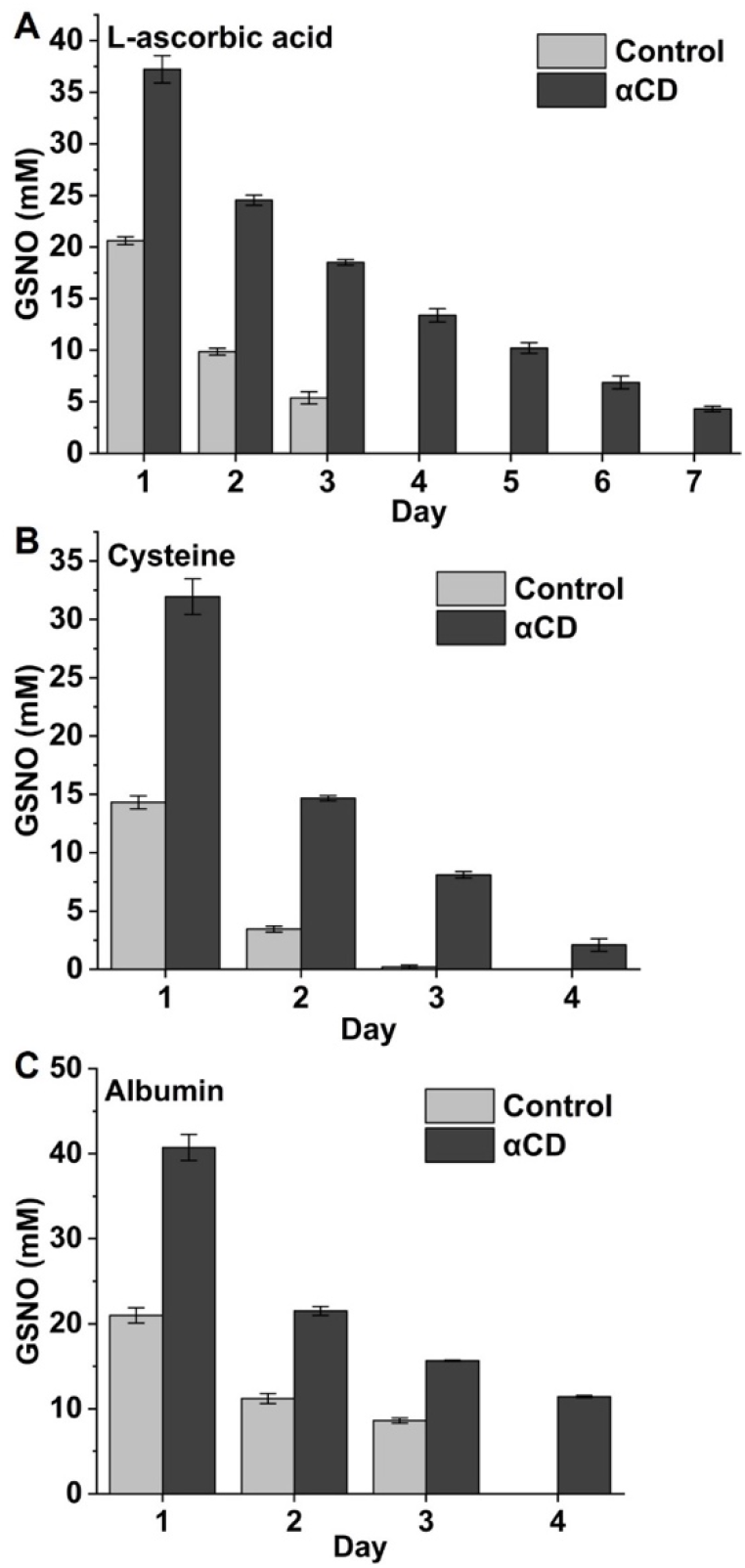
Decomposition of 0.1 M GSNO in 0.1 M PBSE with and without 0.1 M αCD in the presence of 10 mM L-ascorbic acid (A), 10 mM cysteine (B), and 20 mg/mL BSA (C) at pH 7.4 and 37°C.

### 3.4 NO release and antibacterial properties of catheters filled with GSNO-CD solutions

The GSNO-CD solutions with controlled and sustained release of NO have a variety of potential applications. First, catheters such as central line catheters and urinary Foley catheters are always filled with lock solutions or inflation solutions. NO release from these solutions can reduce the infectious and/or thrombotic complications of catheters due to the potent antimicrobial and antiplatelet activities of NO [20–25]. Since one solution may need to fill urinary catheters and central line catheters for days to multiple weeks, sustained NO release is necessitated. Second, we and other groups found that NO reduces inflammation and infection of insulin infusion cannulas and holds promise in enhancing the lifetime of such subcutaneously implanted cannulas [58,59]. The tiny Teflon or stainless-steel cannulas cannot hold a significant amount of NO donors. Instead, concentrated GSNO may be infused into cannulas together with insulin to supply a larger amount of NO. The NO donor needs to undergo minimal degradation in the infusion pump for one week or more. Third, NO-releasing nasal sprays have proven to be highly effective in mitigating viral infections including COVID-19 [60]. The GSNO solution stabilized by CDs may be a formulation that is amenable for storage and releases NO for a prolonged period of time in the respiratory tract. GSNO-CD solutions with sustained NO delivery may also promote other therapeutic applications for diseases such as cancer, stroke, asthma, embolization, and cystic fibrosis [18,29,32,61].

In this paper, we use lock solutions of central line catheters as an example application of GSNO-CD solutions and focus on the antibacterial activities of NO to prevent and treat infections. More than 5 million central venous catheters are inserted annually in the US to provide venous access for kidney dialysis, anti-cancer drug infusion, total parenteral nutrition, long-term antibiotic treatment, and repeated drawing of blood samples. Approximately 30,000 episodes of central line-associated bloodstream infections occur in USA acute care hospitals annually [62]. There are also ∼ 30,000 bloodstream infections in outpatient hemodialysis facilities annually in the US, 76.5% of which are related to hemodialysis catheters [63]. Compared to coating or doping bactericidal agents onto or into polymeric catheters, locking catheters with antimicrobial solutions between uses is a replenishable strategy without limitations in drug loading. Direct use of commercial catheters without any device modification is another advantage of the lock solution method. Antibiotics and ethanol are currently used in clinical applications as anti-infective lock solutions [64]. Other drugs like taurolidine are being developed for catheter locking applications [65]. One obvious drawback of these locking solutions is that the large organic drugs cannot pass through the wall of a catheter and therefore cannot protect the exterior surface of the catheter at all although bacterial contamination may occur extraluminally due to open surgical site contamination, exit-site infection or bacteremia caused by other sources [66]. Due to the high diffusivity of NO through the polymer materials [67], the NO release solution can fully protect the entire catheter (both intraluminal and extraluminal surfaces) from bacterial colonization.

#### 3.4.1 NO release of silicone rubber tubes filled with GSNO-αCD solutions

The NO generation from the solution itself has been tested (Figure 2), which is indicative of the NO generation within the catheter lumen from the GSNO-CD lock. To evaluate the NO release from the exterior catheter surface, we filled medical-grade silicone rubber tubes with GSNO or GSNO-αCD solutions and sealed both ends. As shown in Figure 6A, NO indeed penetrates through the polymeric catheter wall to reach the outer catheter surfaces and be detected by the NO analyzer. The 0.1 M GSNO solution and 0.1 M GSNO + 0.1 M αCD solution release NO over 0.1 × 10^−10^ mol cm^-2^ min^-1^ for 6 days and 11 days, respectively, from the catheter exterior. αCD suppresses the initial release of NO from 14.8 × 10^−10^ mol cm^-2^ min^-1^ to 6.3 × 10^−10^ mol cm^-2^ min^-1^ on day 0. Increasing αCD to 0.15 M further extends the NO release duration and reduces the initial release to 3.9 × 10^−10^ mol cm^-2^ min^-1^, which is comparable to the 0.5 - 4 × 10^−10^ mol cm^-2^ min^-1^ flux of NO released from the healthy endothelium in the human body [68]. The extraluminal NO release from catheters can be further adjusted by changing the concentration of GSNO. For example, a one-week release of NO above a flux of 0.5 × 10^−10^ mol cm^-2^ min^-1^ is obtained by using 0.15 M GSNO + 0.15 M αCD (Figure 6B). In previous studies using nitrite or S-nitrosothiol-based solutions/suspensions as filling solutions of catheters, only less than 2 days of NO release with initial bursts was reported, and the antibacterial efficacy was only assessed with < 24 h of locking using NO donor solutions [20–23,25]. Given that central line catheters need to be locked for 2 or 3 days to several weeks in hemodialysis and chemotherapy and the GSNO reactivity is higher in the real biological environment containing hemoglobin and other reactive species [29,33,57,69], more steady and sustained NO release from the GSNO-CD solutions is desirable.

**Figure 6.**
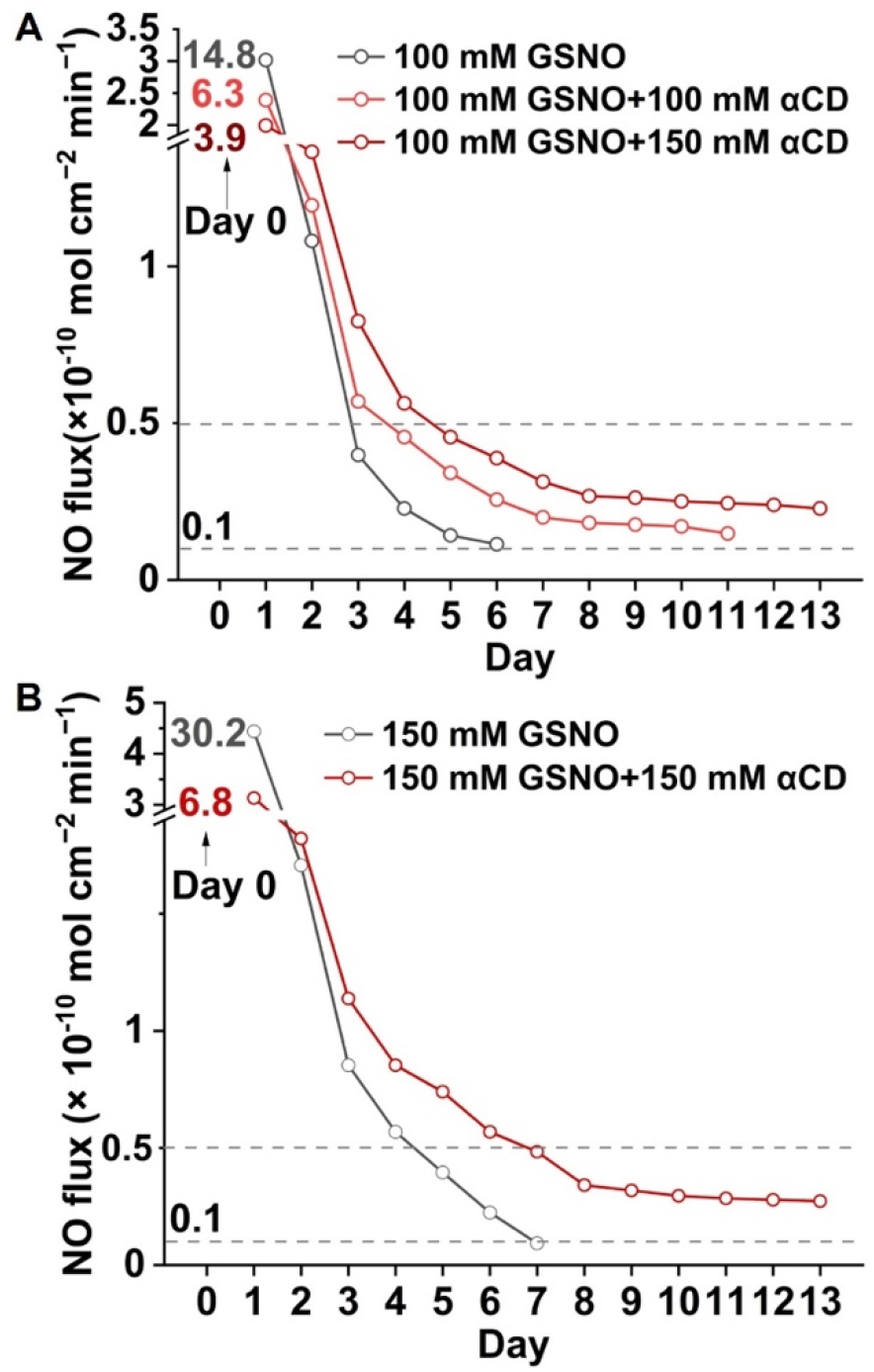
NO release from silicone catheters filled with 0.1 M GSNO (A) and 0.15 M GSNO (B) mixed with different concentrations of αCD in pH 7.4 PBSE at 37°C. The initial NO fluxes on day 0 are shown as numbers.

#### 3.4.2 Efficacy of the GSNO-αCD solution in preventing and treating extraluminal catheter infections

Intravascular catheters are prone to microbial biofilm colonization on intraluminal and extraluminal surfaces [66]. Prevention and treatment of extraluminal biofilms by traditional antimicrobial lock solutions is hardly possible because the drug resides within the catheter lumen and cannot penetrate the catheter wall. Herein, we assess the effectiveness of GSNO-αCD lock solution against extraluminal biofilm formation using *S. aureus*, the most causative microorganism in intravascular catheter-associated infections [70], as an example. The *S. aureus* biofilm on the outer surface of silicone rubber catheters filled with different NO release and control solutions is quantified using the plate counting method. As shown in Figure 7A the GSNO only and GSNO-αCD solutions reduce bacterial biofilm by 70% and 88%, respectively, compared to the control solutions using phosphate buffer and αCD in 3-day experiments. Because of the sustained NO release of GSNO when complexed with αCD, the antibacterial effect is also more sustainable. As shown in Figure 7B, although the GSNO-αCD group shows slightly more bacteria than the GSNO group on day 1 due to the lower initial flux, its biofilm bacteria become significantly less on days 3 and 5 as the NO flux from GSNO-CD surpasses that from GSNO only.

**Figure 7.**
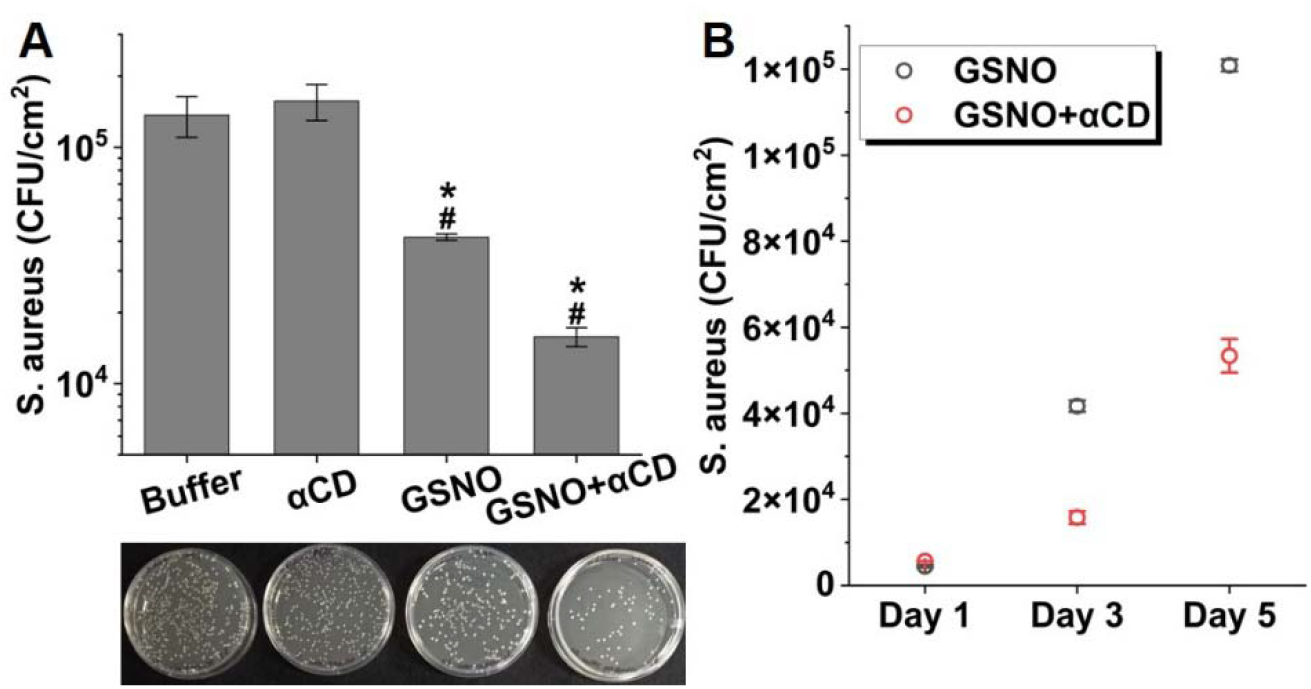
Prevention of *S. aureus* biofilm growth on the extraluminal surface of silicone catheters by GSNO lock solutions. (A) Viable biofilm bacteria on the outer surface of silicone catheters filled with PBSE buffer as the control, or PBSE containing 0.1 M αCD, 0.1 M GSNO, or 0.1M GSNO+0.1 M αCD after 3-day incubation with *S. aureus* at 37°C. (B) Comparison of the biofilm growth on the external catheter surface after incubation with the *S. aureus* culture at 37°C for 1, 3, and 5 days. The catheter is filled with 0.1 M GSNO solution with and without 0.1 M αCD. *: p <0.05 compared to control. ^#^: p <0.05 compared to αCD.

Eradicating established biofilms on the catheter surface is another clinical need because removing a contaminated catheter is not always a first-line option. Current guidelines recommend sterilizing a previously infected catheter with a highly concentrated antibiotic (100–1,000 times planktonic minimum inhibitory concentration) for 24 to 48 h to eliminate biofilms [64]. Again, this method is effective primarily for bacterial biofilms on the intraluminal surface rather than the extraluminal surface. NO is a unique therapeutic agent against mature biofilms because it can easily penetrate biofilms to disperse and/or kill bacteria [71]. Therefore, the NO release from GSNO-αCD solutions is used to remove *S. aureus* biofilms developed on the outer catheter surface. Silicone rubber tube segments with plugged ends were exposed to *S. aureus* culture for 48 h to grow biofilms on the outer surface. Then, these tubes were filled with different lock solutions for 24 h with both ends sealed before the bacterial biofilm was quantified. As shown in Figure 8A, both NO release solutions reduce the number of viable *S. aureus* on the outer surface by at least 90% compared to the control catheters treated with phosphate buffer or αCD. The slightly high efficacy of the GSNO solution without αCD after 24 h agrees with its higher NO flux in the first day. After 48 h, the GSNO-αCD solution becomes more effective due to its higher NO flux on day 2 (Figure 8B). More than 99% of mature *S. aureus* biofilm on the outer surface is removed after the 48-h treatment using the GSNO-CD solution, representing a unique lock therapy for the eradication of extraluminal bacterial biofilms.

**Figure 8.**
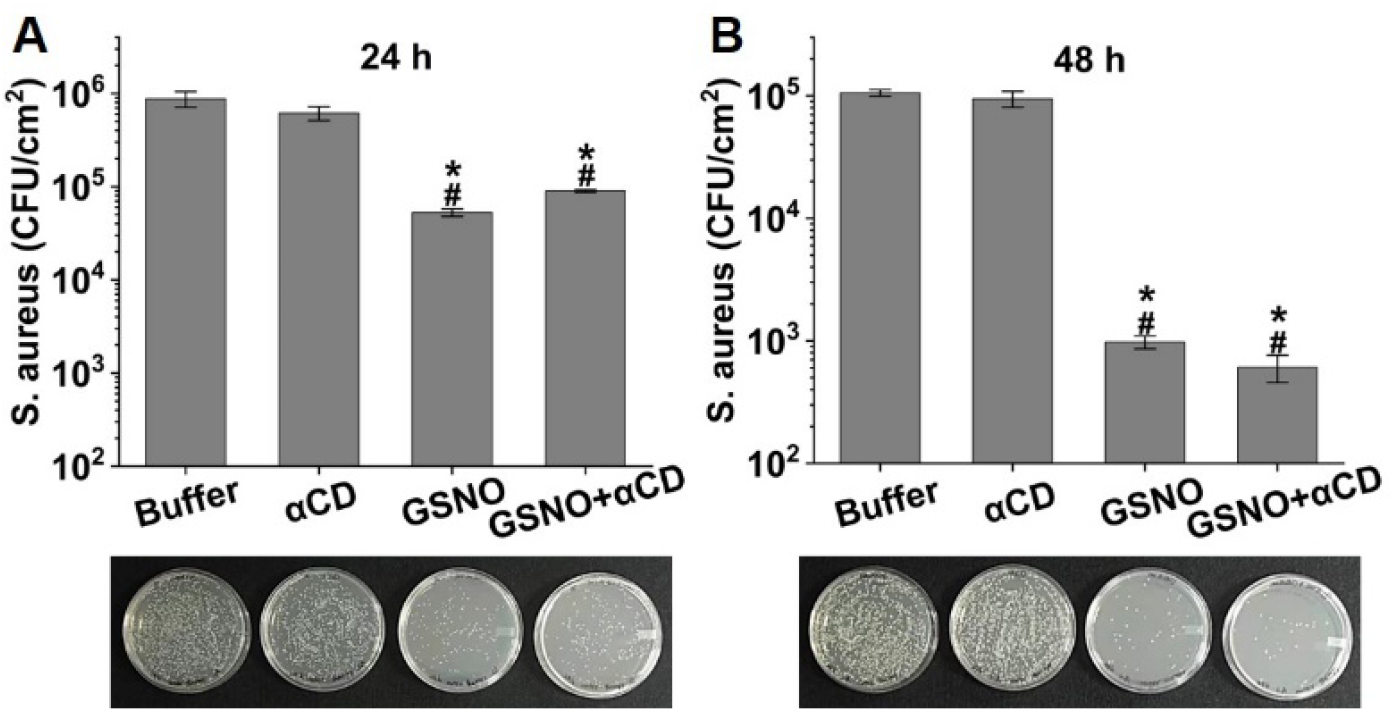
Eradication of mature *S. aureus* biofilm on the extraluminal surface of silicone catheters by lock solutions containing 0.1 M GSNO with and without 0.1 M αCD. The viable bacteria were counted after 24-h (A) and 48-h (B) treatments. *: p <0.05 compared to control. ^#^: p <0.05 compared to αCD.

#### 3.4.3 Efficacy of the GSNO-αCD solution in preventing and treating the intraluminal catheter infections

Bacteria also enter the catheter lumen from hub region contamination and colonize the luminal surface to form biofilms [72]. Unlike exterior surfaces that are only exposed to NO diffused through the catheter wall, the lumen is filled with the lock solution. An antibacterial lock solution should kill planktonic bacteria in the solution and inhibit biofilm development on the intraluminal surface. Therefore, we assessed the growth of both planktonic and biofilm bacteria in the presence of different lock solutions in 3-day experiments. As shown in Figure 9, the growth of planktonic *S. aureus* is reduced by 5 and 6 orders of magnitude by the GSNO and GSNO-αCD solution, respectively, compared to the phosphate buffer. The viable *S. aureus* in the biofilm on silicone rubber is undetectable when the silicone rubber is exposed to the GSNO-αCD solution. The bacterial biofilm is reduced by 99.2% in the GSNO only group but more than the GSNO-αCD group, which again agrees with the higher NO flux generated by the GSNO-αCD inclusion complex after the first day. If the *S. aureus* biofilm is already established on the silicone rubber surface, the use of GSNO-αCD solution completely eradicates the biofilm after one day of lock therapy (Figure S2).

**Figure 9.**
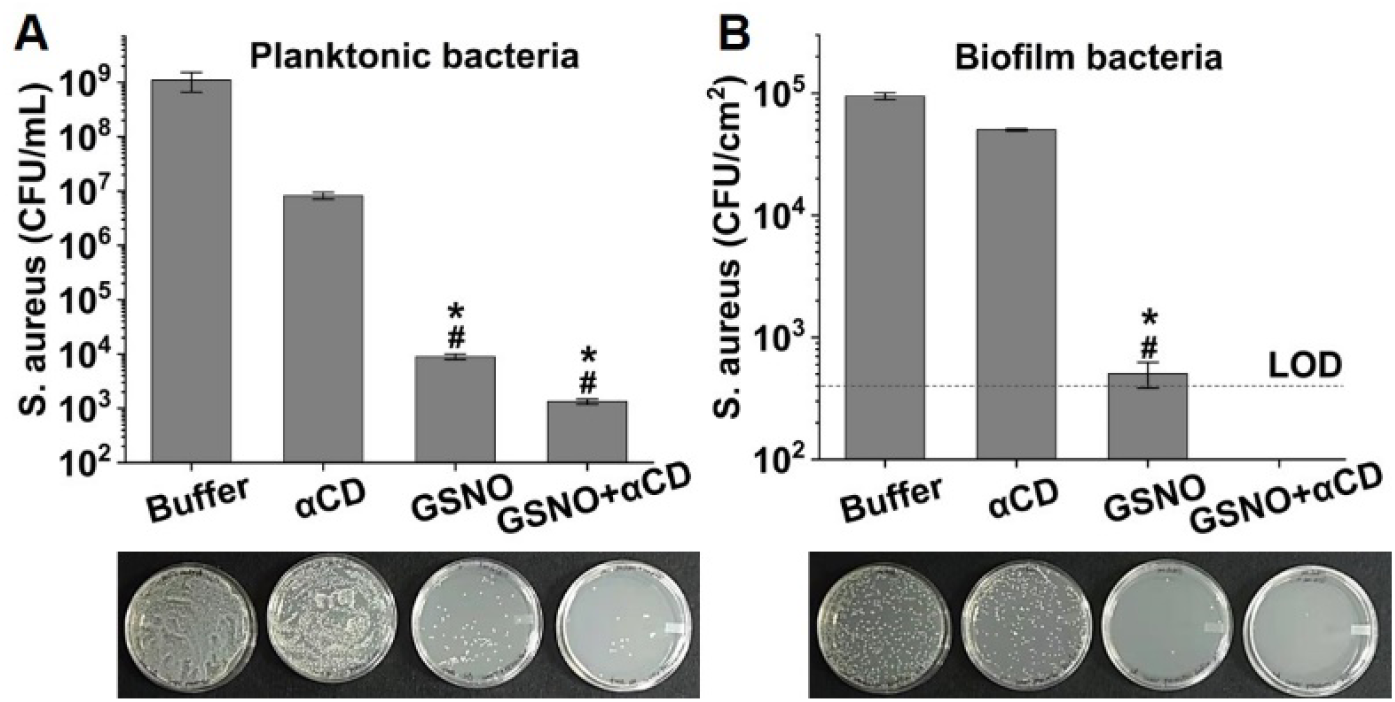
Prevention of intraluminal catheter infection by NO-releasing lock solutions containing 0.1 M GSNO with and without 0.1 M αCD compared to control solutions. Viable planktonic bacteria in lock solutions (A) and viable biofilm bacteria on the intraluminal surface (B) after 3-day incubation with *S. aureus* at 37°C. The dashed gray line at 400 CFU/cm^2^ represents the detection limit of our method. *: p <0.05 compared to control, ^#:^ p <0.05 compared to αCD.

#### 3.4.4 Broad-spectrum activities of NO against multiple bacterial strains

NO is known to exert broad-spectrum antibacterial activities based on multiple nitrosylation and oxidation mechanisms toward enzymes, proteins, DNA, and lipids [73]. To confirm the efficacy of GSNO and GSNO-αCD solution against other bacteria, we tested two more gram-positive strains (MRSA and *S. epidermidis*) and two gram-negative strains (*E. coli* and *P. aeruginosa*). Figure 10 shows the viable planktonic bacteria after 3 days of incubation with different NO release and control solutions. The GSNO-αCD solution results in at least 6-log reductions of all bacterial strains, confirming the highly broad-spectrum antibacterial activities of NO. Similarly, the formation of bacterial biofilm is also inhibited by NO release solutions to the level of being undetectable under our experimental conditions except *E. coli*. The GSNO solution only reduces *E. coli* biofilm by 79% while the GSNO-αCD solution leads to 99% reduction. The reason of the low efficacy of the GSNO solution in preventing *E. coli* biofilm formation on the silicone surface is not fully understood, but it may be related to the reaction of NO donors with indole generated by *E. coli* as demonstrated in our recent publication [74]. This reaction may consume NO donors and compromise the NO release. The much better efficacy of GSNO-αCD solution is an interesting observation, suggesting that αCD probably suppresses such reactions as it does for other GSNO reactions and preserves the NO for bacterial inhibition.

**Figure 10.**
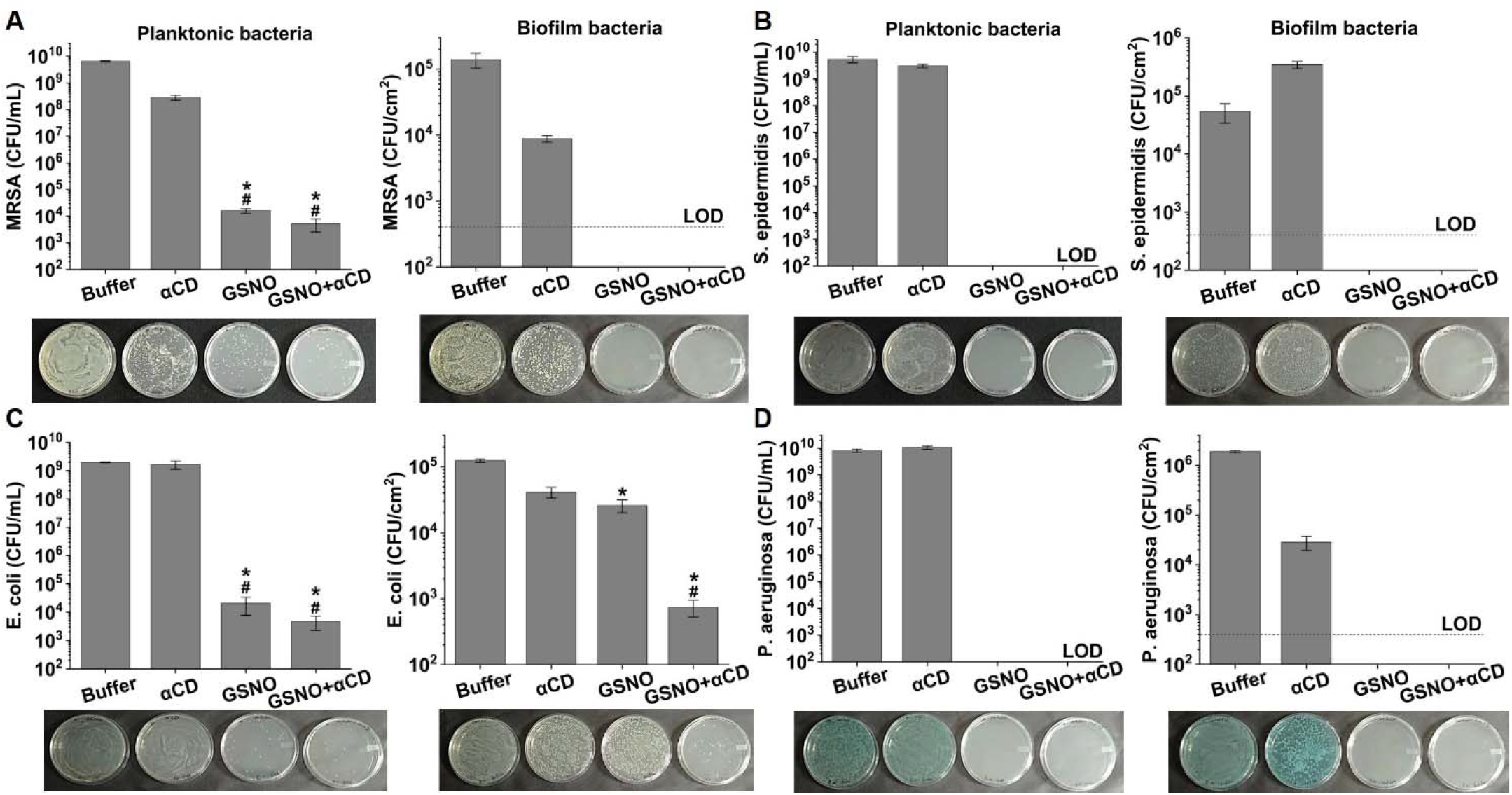
Antibacterial effect of NO release solutions on multiple strains including MRSA (A), *S. epidermidis* (B), *E. coli* (C), and *P. aeruginosa* (D). Planktonic bacteria and biofilm bacteria on the silicone catheter surface were quantified after 3-day exposure to different NO release and control solutions. The dashed gray line represents the detection limit of the method. *p <0.05 compared to control, ^#^p <0.05 compared to αCD.

#### 3.4.5 Cytocompatibility of the NO-releasing catheter

MTS assays were performed to evaluate the possible cytotoxicity of the NO-releasing catheter filled with the GSNO or GSNO-αCD solution as a lock solution. As shown in Figure 11, there is no difference in the cell viability between silicone rubber tubes filled with NO-releasing solutions, phosphate buffer, and αCD, indicating that the NO release induces no cytotoxicity to mouse fibroblast L929 cells.

**Figure 11.**
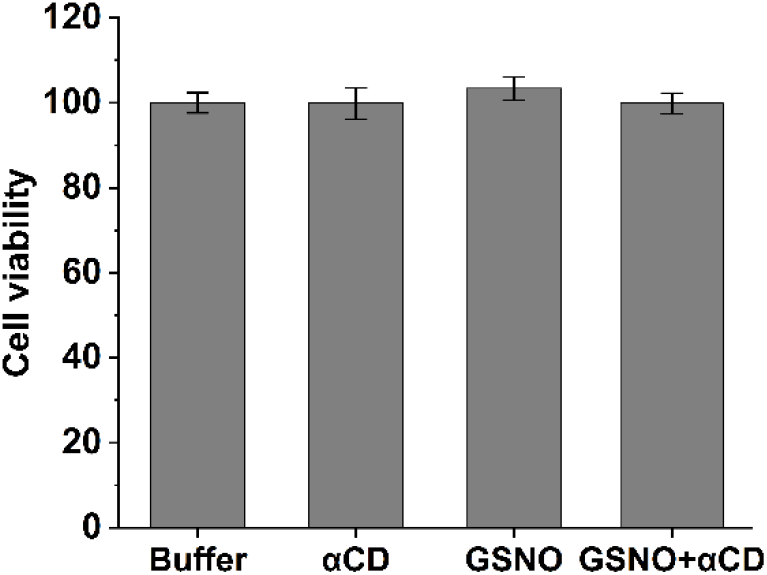
Viability of fibroblast L929 cells after 24-h incubation with silicone catheters locked with phosphate buffer, 0.1 M αCD solution, 0.1 M GSNO solutions with and without 0.1 M αCD.

### 3.5 Antibacterial property of the GSNO-αCD hydrogel

We also tested the antibacterial property of the NO-releasing agar hydrogel using the agar diffusion method. As shown in Figure 12, the agar hydrogel containing GSNO-αCD grows fewest *S. aureus* (less white) and the GSNO hydrogel also grows less bacteria compared to the hydrogel with buffer or αCD. This is consistent with the high antibacterial activity of GSNO-αCD solution.

**Figure 12.**
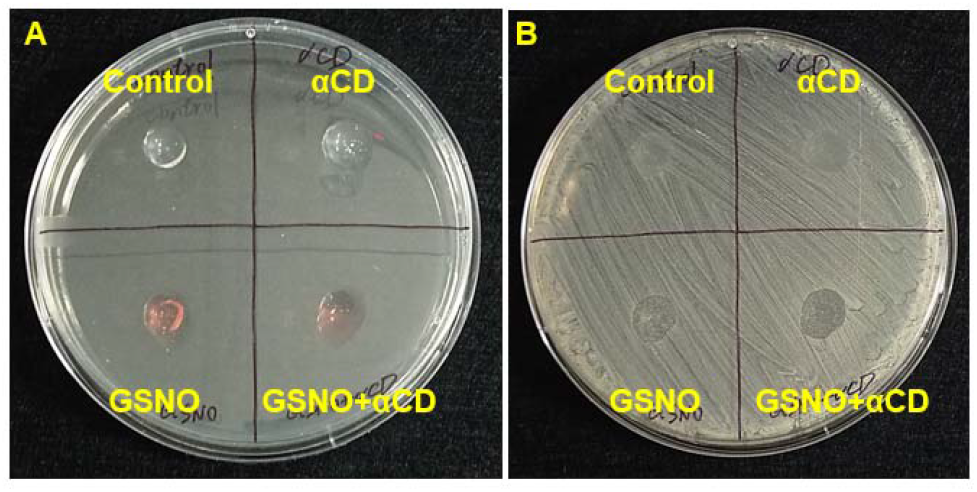
Photos of hydrogels on an LB-agar plate inoculated with *S. aureus* before (A) and after (B) 24-h incubation at 37°C.

## 4. Conclusions

We demonstrated that GSNO at near-neutral pH forms complexes with CDs and becomes chemically more stable. αCD is most effective in stabilizing GSNO due to the highest bond dissociation energy of S-NO after the conformational change of GSNO in the CD cavity. The GSNO-αCD solution releases NO with a significantly reduced initial burst and an extended duration. The sustained NO release from the GSNO-CD solution and hydrogel enables highly effective prevention and treatment of bacterial infections. In addition to thermal decomposition, the reactivity of GSNO in the presence of biological catalysts and light is also regulated by the host-guest complexation. As there are many more CD derivatives and other macrocyclic molecules available, more NO release profiles may be obtained from host-encapsulated GSNO. Our new GSNO formulations are also expected to advance more applications of NO such as thrombosis inhibition, cancer therapy, virus eradication, and wound healing.

## Acknowledgement

X.W. acknowledges JDRF (1-FAC-2019-874-A-N), VCU’s CTSA (UL1TR002649 from the National Center for Advancing Translational Sciences), and the CCTR Endowment Fund of Virginia Commonwealth University. This work is also supported by the start-up fund from the Virginia Commonwealth University (X.W. and K.U.L.). This research used resources of the National Energy Research Scientific Computing Center, which is supported by the Office of Science of the U.S. Department of Energy under Contract No. DE-AC02-05CH11231. High Performance Computing resources provided by the High Performance Research Computing (HPRC) Core Facility at Virginia Commonwealth University (https://chipc.vcu.edu) were also used for conducting the research reported in this work.

