## supporting information for "Inclusion Complexation of S-Nitrosoglutathione for Sustained NO Release and Reduced Device Infection"

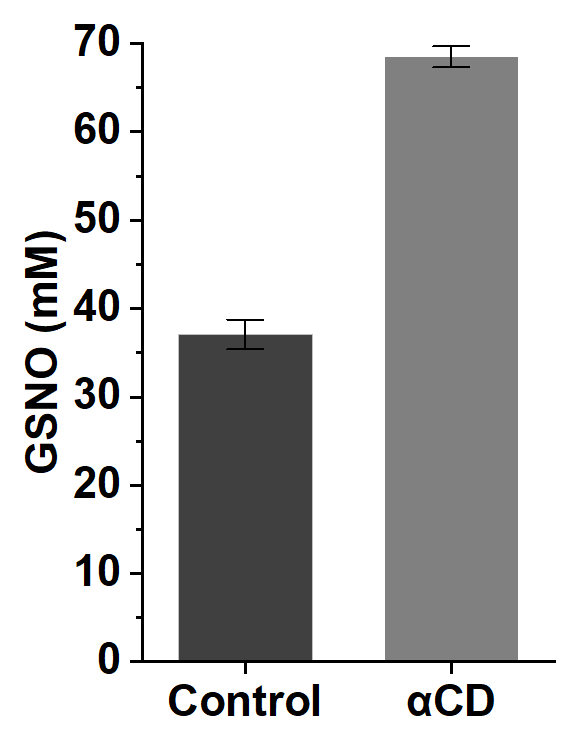

**Figure S1.** Photolytic decomposition of 0.1 M GSNO with and without 0.1 M αCD in PBSE after exposure to white light for 4 h.

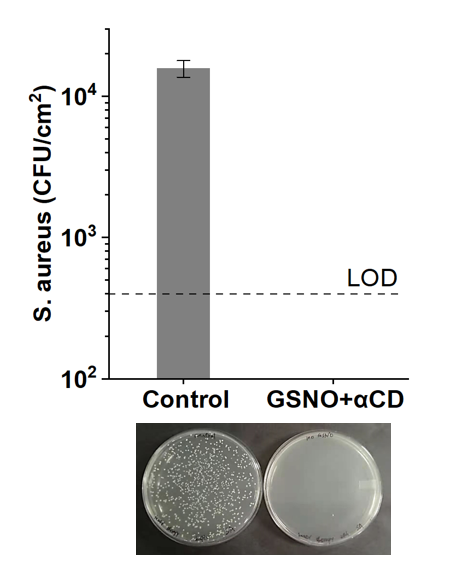

**Figure S2.** Eradication of mature *S. aureus* biofilms on the surface of silicone catheters by exposing the surface to a 0.1 M GSNO + 0.1 M αCD solution for 24 h at 37°C. The control group does not have GSNO or αCD.

**The Cartesian coordinates of the free GSNO molecule, isolated alpha-/hp-beta-/gamma-CDs, and three GSNO-alpha-CD, GSNO-hp-beta-CD, and GSNO-gamma-CD optimized at the level of r2SCAN-3c with CPCM (water)**

(1) The free GSNO molecule

-1 1

N	5.268051	4.868521	-1.661197

H	4.255215	4.694630	-1.823289

H	5.770798	4.237288	-2.310488

C	5.681256	4.588095	-0.248866

C	5.685365	3.091684	0.052497

H	5.874583	2.997295	1.123936

H	4.686314	2.673936	-0.132541

C	6.753750	2.262548	-0.704353

C	6.212723	1.781003	-2.029958

O	6.077155	2.546959	-3.001565

N	5.817594	0.492813	-2.077440

C	5.036202	-0.053872	-3.181750

C	5.412538	-1.500932	-3.470964

H	6.466967	-1.548053	-3.779256

N	7.032037	-2.984818	-1.626250

O	7.862940	-2.461647	-2.281132

C	3.527110	0.105293	-2.877221

O	2.804577	-0.869473	-2.643975

N	3.104398	1.376180	-2.860955

H	3.751204	2.138432	-3.044137

H	7.646197	2.869692	-0.895159

H	7.057174	1.413797	-0.083900

H	4.932333	5.067787	0.394762

C	7.041140	5.291703	0.035602

O	7.433780	6.117658	-0.828698

O	7.589878	4.986882	1.119659

H	5.529092	5.831191	-1.887244

H	4.791393	-1.893154	-4.280790

H	1.526628	1.239256	-1.478831

H	5.859620	-0.054586	-1.226434

C	1.789030	1.782916	-2.394442

H	1.013308	1.551140	-3.137969

C	1.741821	3.298217	-2.109420

O	2.756618	3.980169	-2.446202

O	0.688971	3.720193	-1.586973

S	5.241348	-2.620811	-2.047906

H	5.276352	0.539168	-4.073142

(2) Isolated alpha-CD

0 1

C	-2.461019	4.047550	-0.986244

O	-2.044333	4.907296	-2.050553

H	-2.431092	4.561901	-2.866929

H	-2.111024	3.019515	-1.175630

C	-3.982566	4.032354	-0.844567

O	-4.607025	3.494466	-2.005911

H	-4.600343	2.524125	-1.920602

H	-4.339565	5.065025	-0.738290

C	-4.379057	3.256557	0.412053

O	-3.734837	3.810663	1.544861

C	-2.294293	3.724634	1.489742

C	-1.771602	4.184523	2.853214

H	-0.973725	4.920604	2.734895

O	-1.206243	3.117209	3.623622

H	-1.851878	2.381116	3.688554

H	-2.600558	4.657547	3.397914

H	-1.990366	2.677797	1.337079

C	-1.792613	4.538881	0.291077

O	-0.389162	4.327296	0.102108

C	0.522516	5.337712	0.479744

O	1.151716	5.048699	1.713205

C	1.847875	3.784020	1.734168

C	2.330789	3.577194	3.169405

O	1.549090	2.619367	3.894240

H	0.595822	2.822342	3.781802

H	3.353827	3.195373	3.176956

H	2.321483	4.551004	3.679217

H	1.151772	2.972227	1.473278

C	2.964414	3.800036	0.680653

C	2.384520	4.141148	-0.684929

O	3.474477	4.246732	-1.604395

H	3.097287	4.440736	-2.473680

C	1.566803	5.429802	-0.634092

O	0.960069	5.707701	-1.891766

H	0.140014	5.183558	-1.945501

H	2.236589	6.269900	-0.408716

H	1.721597	3.312426	-0.984338

H	3.742771	4.529006	0.955902

O	3.527578	2.490443	0.543237

C	4.806483	2.218971	1.073425

O	4.735384	1.567966	2.329115

C	3.965184	0.346824	2.303957

C	3.838981	-0.125516	3.754565

O	2.506593	-0.008277	4.266969

H	2.179965	0.905300	4.120109

H	4.100243	-1.182145	3.842803

H	4.541575	0.455815	4.367889

H	2.956345	0.551558	1.914390

C	4.642059	-0.651178	1.357068

O	3.806515	-1.798588	1.170166

C	4.132204	-3.007742	1.822516

O	3.316053	-3.234171	2.955648

C	1.901132	-3.246598	2.668431

C	1.189522	-3.366703	4.013870

O	0.665280	-2.120117	4.488400

H	1.343948	-1.417850	4.394553

H	0.333250	-4.039385	3.931805

H	1.895492	-3.793335	4.740661

C	1.587940	-4.379891	1.680872

H	1.752455	-5.360088	2.155256

C	2.456365	-4.244023	0.438404

O	2.206236	-5.380314	-0.392264

H	2.741580	-5.272097	-1.190562

C	3.934854	-4.130651	0.803404

O	4.739245	-3.928548	-0.353799

H	4.706837	-2.979639	-0.574093

H	4.264552	-5.072258	1.261390

H	2.151011	-3.323025	-0.085731

O	0.235317	-4.267342	1.220822

C	-0.740200	-5.172621	1.688994

O	-1.507313	-4.629047	2.748969

C	-2.138786	-3.373596	2.415747

C	-2.751506	-2.824621	3.707618

O	-2.100272	-1.638222	4.176413

H	-1.137131	-1.804413	4.265505

H	-2.711923	-3.614647	4.470512

H	-3.796846	-2.548638	3.552812

C	-3.136032	-3.611608	1.275964

C	-2.424040	-4.240314	0.086328

O	-3.416064	-4.544782	-0.898424

H	-2.954901	-4.885701	-1.677102

C	-1.648087	-5.490591	0.499637

O	-0.905206	-6.024015	-0.591621

H	-0.067961	-5.529451	-0.642818

H	-2.361430	-6.264785	0.810556

H	-1.714406	-3.497885	-0.314550

H	-3.952476	-4.266710	1.618374

O	-3.657940	-2.364567	0.806020

C	-4.976168	-1.990491	1.148718

O	-4.999448	-1.021336	2.178654

C	-4.256232	0.178020	1.873085

C	-4.280362	1.035084	3.136981

O	-3.049542	0.992049	3.870236

H	-2.756836	0.060552	3.964460

H	-5.118223	0.699696	3.764758

H	-4.441393	2.085061	2.883570

C	-4.859912	0.854410	0.634416

C	-4.919246	-0.132163	-0.523107

O	-5.591121	0.514138	-1.607397

H	-5.594643	-0.104546	-2.350872

C	-5.622227	-1.425888	-0.117402

O	-5.606340	-2.376500	-1.177215

H	-4.744224	-2.831212	-1.154444

H	-6.675708	-1.208478	0.101811

H	-3.882643	-0.370283	-0.814008

H	-5.867455	1.239395	0.857997

O	-4.003514	1.914366	0.192491

H	-3.212389	-0.081166	1.638622

H	-5.542789	-2.849479	1.531769

H	-1.380577	-2.659458	2.059969

H	-0.275312	-6.083279	2.089957

H	1.609729	-2.296577	2.194684

H	5.165193	-2.989495	2.194742

C	4.809122	-0.030570	-0.022959

O	5.543806	-0.957620	-0.827288

H	5.601327	-0.583914	-1.717444

C	5.517404	1.320903	0.059626

O	5.597647	1.943705	-1.218355

H	4.751493	2.399873	-1.375705

H	6.547948	1.163334	0.403591

H	3.803801	0.121181	-0.448932

H	5.623611	-0.950229	1.757175

H	5.368917	3.146143	1.246500

H	0.009655	6.298388	0.621151

H	-2.003709	5.609509	0.439651

H	-5.456474	3.339705	0.607307

(3) Isolated hp-beta-CD

0 1

O	7.798039	-1.523469	2.258104

H	7.700199	-2.084244	3.040704

C	6.871906	-1.996574	1.276547

C	7.006517	-1.117553	0.039753

O	6.646928	0.217896	0.414859

C	7.630599	1.227589	0.278779

O	7.432819	2.014057	-0.879352

C	6.177805	2.721637	-0.928872

C	6.186558	3.505397	-2.222314

H	5.362917	4.229129	-2.223370

H	7.136119	4.053881	-2.289012

C	6.028169	3.638511	0.294856

C	6.236911	2.845987	1.574710

O	6.201090	3.771657	2.665512

H	6.286247	3.258293	3.480710

C	7.571600	2.109058	1.525898

O	7.798937	1.373227	2.721204

H	7.525433	0.448171	2.574287

H	8.376294	2.853485	1.443654

H	5.417941	2.115890	1.674791

O	4.701969	4.175112	0.359568

C	4.494268	5.527783	-0.006987

O	3.788373	5.634523	-1.225706

C	2.486540	5.012625	-1.213682

C	1.955821	5.098681	-2.624549

H	0.864887	4.971708	-2.629860

H	2.205792	6.083338	-3.044517

C	1.605622	5.709006	-0.171167

C	2.289901	5.624829	1.185718

O	1.490972	6.347641	2.126689

H	1.922015	6.270370	2.989479

C	3.703402	6.200412	1.117002

O	4.368860	6.093082	2.368315

H	4.736629	5.191851	2.442795

H	3.632664	7.271740	0.885642

H	2.344376	4.563120	1.478010

O	0.348148	5.033736	-0.051449

C	-0.829694	5.711470	-0.451555

O	-1.342234	5.214289	-1.672080

C	-1.686319	3.814383	-1.659262

C	-2.736716	3.539863	-0.572296

C	-2.251062	4.059436	0.770972

O	-3.312898	3.879075	1.712967

H	-2.988114	4.175214	2.574518

C	-1.863925	5.532008	0.660166

O	-1.407706	6.045467	1.903885

H	-0.449277	5.873668	1.978910

H	-2.757839	6.108636	0.385481

H	-1.371421	3.471715	1.078628

O	-2.949301	2.131753	-0.418215

C	-4.138559	1.562472	-0.940410

O	-3.872470	0.725004	-2.045830

C	-3.003595	-0.390760	-1.759636

H	-2.045658	-0.024951	-1.358828

C	-3.668095	-1.287866	-0.709222

C	-3.958113	-0.458384	0.532655

O	-4.636862	-1.295034	1.473098

H	-4.818123	-0.756284	2.256290

C	-4.808047	0.758876	0.176704

O	-5.073991	1.558497	1.320813

H	-4.304238	2.138891	1.476794

H	-5.779205	0.407377	-0.197278

H	-2.997549	-0.121409	0.956563

O	-2.785142	-2.347200	-0.320344

C	-3.161818	-3.680438	-0.618489

O	-2.430387	-4.212664	-1.705578

C	-1.002506	-4.260716	-1.508115

C	-0.408410	-4.815741	-2.782437

C	-0.682075	-5.136524	-0.288021

C	-1.425387	-4.605406	0.927498

O	-1.170224	-5.493603	2.019587

H	-1.637837	-5.141884	2.790086

C	-2.921148	-4.523010	0.634623

O	-3.643990	-4.038316	1.758144

H	-3.701825	-3.065867	1.695490

H	-3.289274	-5.537925	0.429630

H	-1.039292	-3.600415	1.162761

O	0.716125	-5.088556	0.017826

C	1.498298	-6.253755	-0.180766

O	2.348930	-6.135924	-1.303664

C	3.264973	-5.024073	-1.245005

C	4.051836	-5.055322	-2.535632

C	4.167086	-5.148838	-0.007630

C	3.326215	-5.323570	1.246932

O	4.224751	-5.570673	2.333067

H	3.692821	-5.629843	3.138550

O	1.524630	-6.627315	2.238332

H	0.766008	-6.016692	2.167130

H	2.896585	-7.406873	0.955054

H	2.769912	-4.389008	1.424035

O	4.924693	-3.950728	0.196259

C	6.269560	-3.906965	-0.244635

O	6.423521	-3.014411	-1.328267

C	6.046807	-1.654402	-1.024487

H	5.024693	-1.635574	-0.617474

C	6.046425	-0.899737	-2.331222

H	6.109241	0.180936	-2.150856

C	7.138150	-3.462115	0.935769

O	6.949292	-4.300882	2.068656

H	5.991620	-4.457717	2.170746

H	8.190609	-3.563964	0.638813

H	6.592282	-4.889200	-0.613508

H	4.839670	-6.014661	-0.117727

H	2.703938	-4.078898	-1.179073

H	0.859090	-7.123369	-0.381434

H	-0.994053	-6.174509	-0.484583

H	-0.615123	-3.244817	-1.332243

H	-4.215774	-3.726823	-0.921872

H	-4.606471	-1.701694	-1.110275

C	-2.752169	-1.087141	-3.075392

H	-2.378685	-2.104002	-2.899032

H	-4.812330	2.345154	-1.313322

H	-3.682056	4.041409	-0.834440

C	-2.209952	3.492055	-3.040625

H	-2.707798	2.515044	-3.029515

H	-2.945879	4.258612	-3.319251

H	-0.790119	3.208938	-1.450966

H	-0.625588	6.775375	-0.628429

H	1.450324	6.763537	-0.448512

H	2.582669	3.951203	-0.937287

H	5.452465	6.038547	-0.169095

H	6.767905	4.453619	0.246001

H	5.340455	2.006074	-0.930789

H	8.628150	0.783864	0.164936

H	8.039840	-1.143539	-0.338748

H	5.840976	-1.893800	1.655317

O	-0.490699	-3.815686	-3.806011

O	-1.770382	-0.333735	-3.799044

O	-1.106347	3.454328	-3.955350

O	3.188725	-4.603737	-3.584555

O	2.567629	4.047463	-3.382953

O	6.021119	2.589064	-3.312297

O	4.812572	-1.218440	-2.986752

C	-1.851536	-0.492701	-5.211164

C	-1.401253	3.845280	-5.290722

C	6.648371	2.956445	-4.533792

C	4.693181	-0.679782	-4.297551

C	3.650907	-4.827105	-4.910532

C	-0.732669	-4.297961	-5.128041

C	-0.517377	-0.056373	-5.779488

O	-2.936892	0.236961	-5.748657

H	-2.064104	-1.546018	-5.449411

C	-1.639120	5.337565	-5.432795

O	-2.476734	3.088181	-5.841075

H	-0.510132	3.542016	-5.852032

O	6.209286	4.233611	-4.991685

C	8.162823	2.871073	-4.477896

H	6.252334	2.233281	-5.255965

O	5.554976	-1.338526	-5.206719

H	5.019287	0.371164	-4.291844

O	4.879360	-4.149554	-5.167508

C	-2.144832	-4.826877	-5.309720

O	0.247051	-5.237042	-5.538100

H	-3.310737	3.549434	-5.672771

H	6.787008	4.919969	-4.628619

H	-0.057065	-6.126818	-5.312671

H	0.641385	-5.082054	-2.612472

H	-0.957652	-5.727946	-3.053011

H	6.908951	-1.206690	-2.939981

H	-3.695448	-1.143339	-3.636687

H	-0.572462	-3.411104	-5.752083

H	4.399360	-6.083611	-2.709332

H	4.920791	-4.391151	-2.453886

H	-0.535718	-0.120223	-6.870898

H	0.284822	-0.695413	-5.398823

H	-0.308476	0.978852	-5.485371

H	-1.728446	5.594644	-6.492384

H	-0.792403	5.883899	-5.007621

H	-2.553468	5.655900	-4.920719

H	8.573204	3.034749	-5.478592

H	8.459824	1.876947	-4.132433

H	8.590487	3.619961	-3.803004

H	-2.319049	-5.043957	-6.367853

H	-2.866760	-4.074210	-4.979711

H	-2.310771	-5.745779	-4.737419

H	2.900004	-4.324388	-5.527907

C	2.334839	-6.471461	1.081096

C	3.745325	-6.297329	-5.278722

H	2.818365	-6.807792	-5.002540

H	3.889338	-6.389060	-6.359053

H	4.585376	-6.791589	-4.779202

H	5.621346	-4.724212	-4.932226

C	3.233859	-0.803452	-4.686601

H	2.924278	-1.854682	-4.654313

H	2.604790	-0.230668	-3.998560

H	3.086572	-0.419997	-5.700152

H	5.273099	-2.273316	-5.240398

C	2.533885	4.260482	-4.789818

O	3.498544	5.215084	-5.187220

H	1.562325	4.694991	-5.070589

C	2.744425	2.910422	-5.442665

H	2.762235	3.018478	-6.530665

H	1.937808	2.224999	-5.166492

H	3.696360	2.480101	-5.110023

H	-2.713616	1.186896	-5.696295

H	4.381862	4.813087	-5.070314

(4) Isolated gamma-CD

0 1

O	-2.044235	-7.358176	1.449563

C	-2.594666	-6.361994	0.584916

C	-4.051540	-6.681330	0.244089

O	-4.881205	-6.649184	1.399370

H	-5.195436	-5.734448	1.512361

C	-4.573915	-5.746853	-0.848815

O	-3.697252	-5.749765	-1.960847

C	-2.369042	-5.281606	-1.632028

C	-1.761550	-6.327392	-0.689527

O	-0.416080	-6.011288	-0.321155

C	0.603182	-6.846919	-0.839486

O	1.240825	-6.261364	-1.955947

C	1.886106	-5.002638	-1.655420

C	3.030355	-5.313155	-0.681796

C	2.428226	-5.866698	0.601493

O	3.496353	-6.173600	1.500230

H	3.098788	-6.523342	2.309668

C	1.609004	-7.116884	0.282183

O	0.975030	-7.641414	1.442277

H	0.093573	-7.232197	1.515263

H	2.304118	-7.888364	-0.077605

H	1.773976	-5.096318	1.043275

O	3.784762	-4.146570	-0.341400

C	5.111411	-4.073634	-0.830858

C	5.994840	-3.504171	0.281576

O	5.872720	-4.252670	1.484792

H	4.951751	-4.562605	1.552762

C	5.714359	-2.018579	0.510017

O	6.689738	-1.450541	1.388145

H	6.621820	-1.912695	2.234951

C	5.782051	-1.277352	-0.816888

O	5.500676	0.100345	-0.557652

C	6.316406	1.057201	-1.212633

C	6.806527	2.065759	-0.170181

O	7.450443	1.419692	0.920364

H	6.943092	0.615826	1.138692

C	5.664203	2.962076	0.301813

O	6.148918	4.017509	1.134622

H	6.595155	3.605882	1.887661

C	4.964246	3.589652	-0.894501

O	3.874803	4.360498	-0.377561

C	3.699956	5.673004	-0.870780

O	2.710966	5.727185	-1.881931

C	1.418436	5.252897	-1.439860

C	0.542107	5.079710	-2.685579

O	0.064118	3.734472	-2.837125

H	0.826075	3.145552	-2.991453

H	-0.347618	5.710515	-2.627186

H	1.126854	5.383947	-3.564310

C	0.922678	6.251518	-0.387221

C	1.888342	6.226434	0.790369

O	1.436454	7.181806	1.752788

H	2.043687	7.143417	2.504597

C	3.303035	6.558097	0.312542

O	4.249616	6.469646	1.370910

H	4.557908	5.546839	1.415530

H	3.307273	7.600254	-0.034467

H	1.875642	5.215147	1.230104

O	-0.374281	5.910427	0.110493

C	-1.441345	6.774800	-0.231839

C	-2.329391	6.936793	1.004336

O	-1.582941	7.372895	2.133658

H	-0.691734	6.983582	2.073884

C	-3.102749	5.650845	1.298459

O	-4.074101	5.863869	2.324687

H	-3.598303	6.148914	3.117113

C	-3.830629	5.201648	0.039290

O	-4.550077	4.005061	0.349910

C	-5.918985	3.969281	-0.011021

C	-6.693634	3.310412	1.132854

O	-6.451367	3.959768	2.374554

H	-5.526656	4.266204	2.377991

C	-6.401897	1.811397	1.211957

O	-7.291123	1.172472	2.131954

H	-7.129033	1.553564	3.005789

C	-6.606038	1.181456	-0.157588

O	-6.309119	-0.211796	-0.039321

C	-7.182467	-1.114868	-0.695860

C	-7.598439	-2.195683	0.305727

O	-8.154699	-1.628928	1.485241

H	-7.619585	-0.850452	1.728744

C	-6.426576	-3.119328	0.625973

O	-6.847541	-4.228747	1.422348

H	-7.246531	-3.866489	2.225729

C	-5.821917	-3.662255	-0.659830

O	-4.695984	-4.459649	-0.279897

C	-5.396578	-2.477049	-1.537385

O	-6.572562	-1.690604	-1.829282

H	-4.672272	-1.869800	-0.972968

C	-4.774162	-2.843820	-2.881275

O	-3.631317	-2.040059	-3.202559

H	-3.892386	-1.100783	-3.152199

H	-4.424338	-3.877119	-2.868074

H	-5.547980	-2.755348	-3.657585

H	-6.560027	-4.279122	-1.195944

H	-5.647251	-2.547790	1.158375

H	-8.391084	-2.795590	-0.161826

H	-8.068837	-0.587473	-1.071512

C	-5.691110	1.891697	-1.161243

O	-6.124009	3.269402	-1.221839

H	-4.654962	1.849510	-0.794194

C	-5.702542	1.346136	-2.594469

O	-4.448849	0.753583	-2.956514

H	-3.779823	1.463149	-2.926437

H	-6.456265	0.566577	-2.721867

H	-5.943703	2.176670	-3.272120

H	-7.653472	1.316283	-0.470866

H	-5.356390	1.650564	1.522984

H	-7.763721	3.428965	0.912891

H	-6.296321	4.984567	-0.189337

C	-2.787844	4.987058	-1.064530

C	-3.311883	4.448798	-2.400515

O	-2.800614	3.141500	-2.695709

H	-1.832458	3.217002	-2.787985

H	-4.400034	4.355969	-2.387692

H	-3.031354	5.159715	-3.189719

O	-2.188343	6.276952	-1.324129

H	-2.017060	4.292104	-0.699180

H	-4.532919	5.985804	-0.283287

H	-2.401218	4.855930	1.601836

H	-3.062647	7.724160	0.780776

H	-1.057867	7.750641	-0.556552

H	0.901659	7.258994	-0.830882

H	1.529383	4.265177	-0.967733

H	4.623115	6.040342	-1.338005

C	4.469898	2.465060	-1.814302

C	3.739806	2.913674	-3.078685

O	2.547549	2.157005	-3.328473

H	2.786609	1.211049	-3.361211

H	3.423346	3.954058	-2.990415

H	4.436757	2.837112	-3.925451

O	5.620109	1.708534	-2.250804

H	3.793775	1.816918	-1.235658

H	5.663594	4.240706	-1.441785

H	4.926469	2.355005	0.853583

H	7.561567	2.700939	-0.653317

H	7.172779	0.566887	-1.693268

C	4.771082	-1.911064	-1.779073

O	5.197644	-3.275327	-1.994236

H	3.777550	-1.907591	-1.306864

C	4.637809	-1.252071	-3.157273

O	3.350355	-0.651507	-3.347260

H	2.689776	-1.368893	-3.311293

H	5.369441	-0.453008	-3.292027

H	4.818603	-2.021596	-3.920428

H	6.792358	-1.378981	-1.243654

H	4.704910	-1.892054	0.935497

H	7.038495	-3.599439	-0.049177

H	5.468258	-5.068667	-1.126793

H	3.691319	-6.065568	-1.139769

C	2.287611	-4.367724	-2.991757

H	3.370390	-4.241934	-3.057161

H	1.966127	-5.039098	-3.799823

O	1.721963	-3.061774	-3.166464

H	0.751155	-3.159306	-3.172834

H	1.167583	-4.331389	-1.161481

H	0.177194	-7.790618	-1.203746

H	-1.792890	-7.311532	-1.182469

H	-2.439912	-4.320489	-1.100790

C	-1.628853	-5.029625	-2.950669

O	-1.118102	-3.690898	-3.039873

H	-1.869820	-3.071943	-3.080922

H	-0.764480	-5.690026	-3.049187

H	-2.320393	-5.234731	-3.778923

H	-5.541230	-6.097988	-1.231329

H	-4.088983	-7.706417	-0.148717

H	-2.536966	-5.372101	1.067375

H	-2.571609	-7.359146	2.260343

(5) GSNO-alpha-CD

-1 1

C	-2.469447	3.900086	-1.052283

O	-1.955145	4.710424	-2.115883

H	-2.339333	4.382037	-2.940926

H	-2.162757	2.852640	-1.202832

C	-3.995695	3.969800	-0.996777

O	-4.578892	3.488544	-2.191995

H	-4.138261	2.630971	-2.436747

H	-4.294079	5.022723	-0.891574

C	-4.503692	3.240126	0.255421

O	-3.881802	3.819392	1.406052

C	-2.452195	3.624446	1.424322

C	-1.933503	4.048911	2.805276

H	-1.234394	4.884981	2.719604

O	-1.219946	3.000186	3.470634

H	-1.842960	2.259921	3.639049

H	-2.790139	4.377327	3.410493

H	-2.225561	2.555495	1.294885

C	-1.847172	4.388860	0.244117

O	-0.443052	4.126992	0.146957

C	0.470067	5.157528	0.474793

O	1.103086	4.912458	1.713883

C	1.841531	3.674340	1.760277

C	2.345631	3.525669	3.192650

O	1.563867	2.603623	3.965752

H	0.614173	2.781915	3.808059

H	3.365548	3.135305	3.202040

H	2.350566	4.519079	3.663290

H	1.169785	2.831108	1.534000

C	2.939666	3.683441	0.687150

C	2.313767	3.929537	-0.681831

O	3.343694	3.968361	-1.667521

H	2.951277	3.597899	-2.483415

C	1.501225	5.225804	-0.655789

O	0.872729	5.504197	-1.898189

H	0.047567	4.985339	-1.947383

H	2.180517	6.064039	-0.449147

H	1.635860	3.087202	-0.893618

H	3.694051	4.455007	0.909543

O	3.550313	2.393615	0.609641

C	4.845052	2.194006	1.131565

O	4.810726	1.545596	2.391181

C	4.076395	0.303679	2.370325

C	3.956963	-0.167860	3.819493

O	2.636711	-0.007138	4.355692

H	2.312243	0.902623	4.188150

H	4.179027	-1.234048	3.897535

H	4.689572	0.383475	4.425399

H	3.062265	0.481131	1.980152

C	4.774167	-0.679480	1.421533

O	3.944014	-1.826632	1.215201

C	4.228042	-3.039955	1.874238

O	3.374692	-3.252397	2.985133

C	1.970401	-3.203212	2.654246

C	1.198342	-3.297496	3.968415

O	0.679439	-2.036591	4.409860

H	1.389530	-1.360502	4.376951

H	0.333987	-3.954992	3.849912

H	1.860648	-3.734313	4.729738

C	1.642169	-4.313169	1.646758

H	1.750603	-5.303405	2.116657

C	2.553222	-4.201502	0.432414

O	2.280497	-5.317749	-0.418823

H	2.825768	-5.212368	-1.210772

C	4.024342	-4.147019	0.838322

O	4.864281	-3.952080	-0.295609

H	4.850763	-3.000758	-0.508855

H	4.312553	-5.105126	1.289699

H	2.300491	-3.263815	-0.090534

O	0.307271	-4.131437	1.159211

C	-0.697238	-5.063997	1.505384

O	-1.467870	-4.627048	2.610933

C	-2.084308	-3.337619	2.397187

C	-2.667699	-2.876092	3.737965

O	-2.141863	-1.611247	4.156678

H	-1.168454	-1.690158	4.232894

H	-2.456187	-3.648580	4.490370

H	-3.750956	-2.747135	3.675331

C	-3.080600	-3.474169	1.242725

C	-2.328819	-3.920608	-0.010705

O	-3.229248	-4.077555	-1.103690

H	-2.932114	-3.444261	-1.787484

C	-1.574320	-5.224297	0.259434

O	-0.795029	-5.642920	-0.853897

H	0.078876	-5.220895	-0.779261

H	-2.306720	-6.018876	0.456204

H	-1.599500	-3.130552	-0.246850

H	-3.858888	-4.209572	1.501629

O	-3.669429	-2.213718	0.922466

C	-4.986673	-1.935815	1.368337

O	-4.988947	-0.931427	2.357322

C	-4.348015	0.291000	1.933789

C	-4.339917	1.192134	3.157381

O	-3.166033	1.019613	3.968874

H	-2.922061	0.071361	3.999263

H	-5.246740	0.988243	3.745571

H	-4.356075	2.240718	2.854501

C	-5.054936	0.882821	0.704510

C	-5.263697	-0.165623	-0.396740

O	-6.154056	0.350775	-1.377131

H	-5.689018	0.394597	-2.246876

C	-5.802241	-1.486096	0.156677

O	-5.822401	-2.510245	-0.832439

H	-4.916843	-2.867632	-0.902582

H	-6.840622	-1.338731	0.482712

H	-4.275566	-0.364478	-0.842485

H	-6.025054	1.322678	0.984972

O	-4.202845	1.878710	0.114000

H	-3.311805	0.076451	1.644249

H	-5.436046	-2.824371	1.831460

H	-1.315669	-2.609188	2.096228

H	-0.255813	-6.022607	1.808440

H	1.735868	-2.239031	2.178185

H	5.251350	-3.047568	2.273006

C	4.935810	-0.049856	0.043013

O	5.697181	-0.955936	-0.763454

H	5.672712	-0.619025	-1.669849

C	5.592210	1.326149	0.117672

O	5.623969	1.950920	-1.163484

H	4.766548	2.402451	-1.290373

H	6.632507	1.217788	0.451459

H	3.930928	0.069438	-0.387253

H	5.754244	-0.977881	1.826394

H	5.359110	3.150143	1.297305

H	-0.049808	6.117865	0.588263

H	-2.027889	5.468324	0.366073

H	-5.581416	3.397883	0.397079

N	4.197313	1.297908	-4.291805

H	5.072753	0.996576	-4.723318

H	4.016146	0.739196	-3.438926

C	3.022517	1.146552	-5.218344

C	2.907431	-0.284028	-5.738823

H	1.941333	-0.346615	-6.246058

H	3.684497	-0.458843	-6.491477

C	3.017660	-1.407135	-4.685445

C	2.188907	-1.117346	-3.453786

O	2.634738	-0.360974	-2.573565

N	0.968902	-1.685911	-3.421057

C	-0.111250	-1.516221	-2.451358

C	-0.025862	-0.263417	-1.562631

H	-1.015778	-0.108564	-1.105137

N	0.099297	-0.047007	1.280139

O	-1.041026	0.251022	1.104932

C	-1.464518	-1.410400	-3.176953

O	-2.461342	-2.000465	-2.730262

N	-1.517006	-0.602834	-4.238136

H	-0.703443	-0.034277	-4.479681

H	2.725219	-2.350731	-5.155974

H	4.055039	-1.514614	-4.349532

H	3.215140	1.811187	-6.070524

C	1.760009	1.723238	-4.512966

O	1.960106	2.634123	-3.667315

O	0.654664	1.282046	-4.901644

H	4.291227	2.275228	-4.010503

H	0.229358	0.625920	-2.147573

H	-3.334096	-1.187725	-5.113434

H	0.762032	-2.331199	-4.171390

C	-2.815025	-0.273670	-4.811795

H	-2.647381	0.342319	-5.702129

C	-3.704031	0.523256	-3.827006

O	-3.125875	1.387154	-3.114091

O	-4.937182	0.264238	-3.835864

S	1.130772	-0.425134	-0.174150

H	-0.176467	-2.409479	-1.816146

(6) GSNO-hp-beta-CD

-1 1

O	7.051552	-1.727711	2.565673

H	6.549279	-2.071347	3.335494

C	6.358175	-2.181550	1.407116

C	6.641247	-1.249248	0.228850

O	6.224940	0.075650	0.576976

C	7.212264	1.090276	0.610065

O	7.157381	1.929710	-0.527197

C	5.927906	2.663078	-0.690551

C	6.089103	3.486517	-1.948238

H	5.292276	4.238077	-2.001659

H	7.060584	3.998846	-1.905506

C	5.655885	3.539117	0.543128

C	5.691804	2.686166	1.800079

O	5.539189	3.559518	2.923673

H	5.485862	3.004856	3.714215

C	7.006443	1.915727	1.881458

O	7.054956	1.115646	3.055254

H	6.837572	0.192241	2.812078

H	7.830666	2.641588	1.939688

H	4.858944	1.967496	1.769633

O	4.346851	4.120969	0.474970

C	4.230818	5.488680	0.108798

O	3.598854	5.635683	-1.145254

C	2.258709	5.104999	-1.194678

C	1.815836	5.160191	-2.635375

H	0.720072	5.118196	-2.699208

H	2.169216	6.095003	-3.092750

C	1.362218	5.877064	-0.223138

C	1.966457	5.755989	1.168766

O	1.172000	6.533704	2.068246

H	1.541173	6.414106	2.954717

C	3.423277	6.212609	1.190245

O	3.995599	6.038337	2.479807

H	4.263858	5.103804	2.571377

H	3.457949	7.288201	0.970009

H	1.926136	4.692003	1.455767

O	0.074008	5.261162	-0.163671

C	-1.066187	5.942833	-0.644914

O	-1.530735	5.411485	-1.873000

C	-1.877192	4.010061	-1.833786

C	-2.966399	3.758241	-0.779018

C	-2.537424	4.328047	0.562734

O	-3.635980	4.172679	1.471198

H	-3.332852	4.467936	2.340866

C	-2.151033	5.798768	0.424073

O	-1.740719	6.344379	1.671776

H	-0.795149	6.136953	1.800268

H	-3.030890	6.370590	0.099662

H	-1.667943	3.753875	0.914946

O	-3.158977	2.348348	-0.591790

C	-4.336522	1.741553	-1.094745

O	-4.057838	0.864624	-2.168760

C	-3.183795	-0.236629	-1.842388

H	-2.224917	0.150163	-1.465946

C	-3.826094	-1.096016	-0.748294

C	-4.117132	-0.213488	0.455278

O	-4.768583	-1.015023	1.444411

H	-4.963214	-0.437566	2.196481

C	-4.995919	0.967796	0.049636

O	-5.287585	1.801591	1.161776

H	-4.541543	2.419732	1.292217

H	-5.955836	0.577850	-0.315620

H	-3.158859	0.162916	0.850622

O	-2.914212	-2.123828	-0.333579

C	-3.334280	-3.473219	-0.448304

O	-2.734975	-4.131751	-1.549003

C	-1.297764	-4.224835	-1.487351

C	-0.832399	-4.930804	-2.739906

C	-0.884850	-5.002725	-0.228123

C	-1.454644	-4.299856	0.993261

O	-1.116614	-5.065463	2.152554

H	-1.480498	-4.604160	2.921107

C	-2.971863	-4.193414	0.851427

O	-3.553333	-3.584935	1.996284

H	-3.704469	-2.638459	1.809974

H	-3.378153	-5.213096	0.785249

H	-1.012951	-3.292300	1.065230

O	0.539822	-5.043819	-0.095450

C	1.209552	-6.282103	-0.311869

O	2.055056	-6.223123	-1.442613

C	3.062947	-5.195608	-1.382344

C	3.940134	-5.366651	-2.602695

C	3.914108	-5.314442	-0.106056

C	3.066891	-5.507280	1.147964

O	3.943587	-5.807810	2.234959

H	3.731121	-5.245290	3.009149

O	1.187278	-6.764028	2.076556

H	0.559667	-6.016054	2.105037

H	2.532224	-7.564725	0.784579

H	2.531796	-4.569892	1.340097

O	4.636070	-4.095486	0.098659

C	6.013314	-4.027569	-0.237401

O	6.230188	-3.104844	-1.283221

C	5.832947	-1.752269	-0.974192

H	4.764370	-1.731211	-0.705997

C	6.016014	-0.952579	-2.242150

H	6.019569	0.122085	-2.018807

C	6.774200	-3.605441	1.021374

O	6.580803	-4.543355	2.067195

H	5.635929	-4.795943	2.083243

H	7.848735	-3.605598	0.792142

H	6.374609	-4.996794	-0.605876

H	4.609752	-6.164786	-0.194591

H	2.586101	-4.204936	-1.394401

H	0.486287	-7.080528	-0.521113

H	-1.290367	-6.025602	-0.282090

H	-0.859823	-3.214835	-1.452528

H	-4.415355	-3.525995	-0.630501

H	-4.760474	-1.543389	-1.121360

C	-2.938000	-0.991958	-3.127378

H	-2.589387	-2.009343	-2.905275

H	-5.020993	2.498948	-1.498967

H	-3.910867	4.231023	-1.094009

C	-2.363766	3.647551	-3.217906

H	-2.885113	2.682792	-3.183369

H	-3.074576	4.418712	-3.547512

H	-0.990968	3.412704	-1.571410

H	-0.840680	6.998885	-0.844221

H	1.279746	6.934192	-0.521251

H	2.266527	4.052628	-0.875552

H	5.224255	5.942709	0.000823

H	6.420050	4.329272	0.617460

H	5.086689	1.963421	-0.816906

H	8.217950	0.650083	0.596774

H	7.715714	-1.257765	-0.012268

H	5.274979	-2.160643	1.602285

O	-1.169827	-4.147950	-3.886067

O	-1.935303	-0.287884	-3.868509

O	-1.237046	3.551418	-4.098849

O	3.233341	-4.943009	-3.775477

O	2.383140	4.021362	-3.295075

O	6.001053	2.605925	-3.075322

O	4.907969	-1.275267	-3.090176

C	-1.912386	-0.605019	-5.254634

C	-1.525506	3.758751	-5.478371

C	6.572664	3.094494	-4.283233

C	5.032167	-0.822363	-4.429382

C	3.936823	-5.124490	-5.011960

C	-0.275970	-4.283732	-4.996766

C	-0.558538	-0.179910	-5.785325

O	-2.980254	0.024172	-5.938235

H	-2.075945	-1.685281	-5.387377

C	-1.790205	5.216890	-5.807946

O	-2.579464	2.916995	-5.936028

H	-0.623364	3.405332	-5.990478

O	6.004779	4.340896	-4.674656

C	8.089274	3.158483	-4.243564

H	6.240479	2.373327	-5.038268

O	6.000127	-1.572873	-5.141949

H	5.403646	0.213809	-4.436306

O	5.149813	-4.388670	-5.031274

C	-0.034250	-5.730323	-5.394487

O	0.931202	-3.585021	-4.777783

H	-3.425813	3.370358	-5.815955

H	6.486706	5.063170	-4.247042

H	1.554771	-4.136217	-4.270517

H	0.252720	-5.068899	-2.652676

H	-1.304809	-5.924255	-2.799827

H	6.970240	-1.222096	-2.717187

H	-3.877272	-1.052535	-3.695611

H	-0.789632	-3.743399	-5.800012

H	4.228998	-6.424440	-2.676430

H	4.841697	-4.753576	-2.481336

H	-0.503354	-0.371038	-6.860565

H	0.240936	-0.732190	-5.282516

H	-0.408054	0.890744	-5.604705

H	-1.883484	5.336883	-6.891234

H	-0.953824	5.827530	-5.455675

H	-2.710638	5.579392	-5.337927

H	8.468312	3.383336	-5.244640

H	8.488771	2.191655	-3.924331

H	8.449249	3.933012	-3.558319

H	0.523062	-5.755268	-6.335580

H	-0.985535	-6.252814	-5.534350

H	0.552828	-6.261087	-4.636855

H	3.279952	-4.644427	-5.745445

C	2.026470	-6.603105	0.944305

C	4.159995	-6.583485	-5.360184

H	3.221551	-7.136535	-5.262425

H	4.505648	-6.655634	-6.395211

H	4.913429	-7.045719	-4.713613

H	5.874571	-4.952066	-4.725736

C	3.653950	-0.924246	-5.050509

H	3.312861	-1.966063	-5.031353

H	2.937219	-0.312516	-4.494315

H	3.685589	-0.578078	-6.087418

H	5.670380	-2.489358	-5.186927

C	2.336587	4.099655	-4.715167

O	3.269208	5.039169	-5.209963

H	1.350946	4.478048	-5.026423

C	2.581680	2.698815	-5.235388

H	2.565221	2.696332	-6.328767

H	1.809899	2.016302	-4.865725

H	3.559182	2.337805	-4.892929

H	-2.804832	0.984393	-5.920949

H	4.165702	4.709468	-5.001655

N	3.384159	-0.663654	5.181913

H	2.614155	-0.017736	5.417792

H	3.744288	-0.347122	4.260851

C	2.933407	-2.083529	5.026439

C	1.811556	-2.201487	3.988309

H	1.328305	-3.165786	4.167559

H	1.054152	-1.429139	4.174497

C	2.240284	-2.185512	2.510969

C	2.634150	-0.832877	1.978299

O	3.510181	-0.133585	2.521417

N	2.006916	-0.423449	0.860523

C	2.215918	0.912214	0.311942

C	2.200012	0.903836	-1.208592

H	2.965383	0.200315	-1.570623

N	0.943124	-1.364248	-2.402781

O	1.981194	-1.829129	-2.069779

C	1.169959	1.850340	0.949598

O	0.173018	2.228118	0.329605

N	1.417682	2.147155	2.236768

H	2.206288	1.709133	2.707706

H	3.115912	-2.833203	2.378585

H	1.431246	-2.607569	1.905891

H	2.541751	-2.402675	5.999144

C	4.145952	-2.979929	4.672923

O	5.270610	-2.421235	4.607292

O	3.874490	-4.193886	4.491702

H	4.125998	-0.592275	5.877203

H	2.438377	1.899605	-1.597531

H	-0.580835	2.395584	2.828371

H	1.216767	-0.956541	0.519827

C	0.422944	2.717391	3.131991

H	0.428452	3.816642	3.097534

C	0.668778	2.283465	4.595262

O	1.696799	1.582157	4.824770

O	-0.179033	2.683291	5.423558

S	0.626859	0.410140	-1.977309

H	3.215159	1.230127	0.628675

(7) GSNO-gamma-CD

-1 1

O	-0.315001	-7.388522	1.848726

C	-1.055628	-6.525684	0.984064

C	-2.462829	-7.067431	0.747152

O	-3.204535	-7.144883	1.956896

H	-3.598130	-6.269175	2.131601

C	-3.197234	-6.247406	-0.317916

O	-2.415599	-6.122948	-1.482979

C	-1.119334	-5.519154	-1.277532

C	-0.312634	-6.428139	-0.339544

O	0.982255	-5.884197	-0.062128

C	2.102972	-6.559990	-0.606680

O	2.532780	-5.967482	-1.814364

C	2.922908	-4.582542	-1.652705

C	4.209136	-4.600107	-0.813254

C	3.863582	-5.152064	0.579779

O	5.016596	-5.289561	1.392557

H	5.017513	-4.550146	2.042364

C	3.219310	-6.537450	0.439019

O	2.752067	-7.048684	1.681170

H	1.817365	-6.794425	1.772104

H	4.003621	-7.222428	0.086527

H	3.142796	-4.463487	1.050699

O	4.794325	-3.302445	-0.647583

C	5.882628	-2.987999	-1.496242

C	7.012647	-2.327822	-0.692413

O	7.474115	-3.146898	0.367713

H	6.926728	-2.967176	1.158901

C	6.562558	-0.951013	-0.217843

O	7.591674	-0.270319	0.500850

H	7.600867	-0.657978	1.390089

C	6.126626	-0.097436	-1.395531

O	5.620757	1.119993	-0.847676

C	6.147852	2.346530	-1.298301

C	6.196000	3.265427	-0.069910

O	6.946500	2.664150	0.977502

H	6.865991	1.697272	0.881938

C	4.775294	3.644344	0.368263

O	4.804236	4.587781	1.436832

H	4.543838	4.092647	2.243070

C	4.042978	4.222181	-0.843706

O	2.747989	4.662620	-0.432745

C	2.373736	5.971616	-0.828039

O	1.485499	5.952116	-1.926024

C	0.245585	5.270137	-1.639163

C	-0.487909	5.067442	-2.969829

O	-0.818975	3.691619	-3.200155

H	0.024114	3.202099	-3.228634

H	-1.431757	5.615546	-2.993416

H	0.156149	5.446644	-3.775176

C	-0.488405	6.097222	-0.578622

C	0.357919	6.104493	0.684607

O	-0.310934	6.893597	1.671869

H	0.211118	6.844387	2.484583

C	1.741222	6.674002	0.375799

O	2.594250	6.625033	1.513045

H	3.115423	5.799501	1.463083

H	1.615693	7.732964	0.108766

H	0.459304	5.067072	1.038874

O	-1.760662	5.536703	-0.243661

C	-2.909569	6.186851	-0.758608

C	-3.899205	6.390736	0.392125

O	-3.296634	7.072146	1.484706

H	-2.392019	6.724471	1.596545

C	-4.512524	5.061878	0.827500

O	-5.565606	5.266910	1.771273

H	-5.185051	5.732080	2.529268

C	-5.090473	4.335870	-0.378246

O	-5.596345	3.084426	0.093219

C	-6.893902	2.697789	-0.315220

C	-7.593001	2.090694	0.902462

O	-7.601404	2.990408	2.004309

H	-6.752110	3.468359	2.006578

C	-6.957099	0.751193	1.272259

O	-7.714125	0.084519	2.284730

H	-7.730777	0.663577	3.059360

C	-6.904331	-0.153530	0.049406

O	-6.264545	-1.370659	0.441979

C	-6.955188	-2.577353	0.180584

C	-6.745958	-3.497194	1.384120

O	-7.167748	-2.883253	2.595499

H	-7.005590	-1.925422	2.520547

C	-5.292630	-3.966062	1.461535

O	-5.141245	-4.976714	2.462033

H	-5.348379	-4.575389	3.316975

C	-4.883603	-4.560929	0.122078

O	-3.520072	-4.977802	0.227368

C	-5.085939	-3.505792	-0.970815

O	-6.497565	-3.195758	-1.006698

H	-4.520411	-2.600609	-0.703698

C	-4.674275	-3.913980	-2.390998

O	-3.494179	-3.230714	-2.829957

H	-3.692465	-2.276480	-2.823137

H	-4.453410	-4.982029	-2.456795

H	-5.516303	-3.694154	-3.062110

H	-5.524009	-5.429446	-0.099429

H	-4.635577	-3.110038	1.687038

H	-7.377415	-4.384153	1.235988

H	-8.024639	-2.386509	0.022883

C	-6.119019	0.563580	-1.053606

O	-6.852844	1.766058	-1.379784

H	-5.126915	0.833006	-0.661212

C	-5.912329	-0.218903	-2.354870

O	-4.525495	-0.449696	-2.643036

H	-4.084930	0.411429	-2.761329

H	-6.375826	-1.206532	-2.294075

H	-6.389506	0.339017	-3.172187

H	-7.924196	-0.361723	-0.309562

H	-5.924743	0.919009	1.623558

H	-8.641593	1.908415	0.630813

H	-7.456192	3.560245	-0.695971

C	-3.965864	4.137509	-1.401681

C	-4.341795	3.380707	-2.674277

O	-3.408028	2.338855	-2.992236

H	-2.522190	2.741263	-3.072812

H	-5.310230	2.891013	-2.557980

H	-4.416523	4.103161	-3.499333

O	-3.499972	5.443730	-1.803310

H	-3.149273	3.588344	-0.909131

H	-5.902485	4.930014	-0.825635

H	-3.728355	4.422584	1.268932

H	-4.708965	7.035274	0.023681

H	-2.638038	7.154497	-1.200081

H	-0.617035	7.126681	-0.948204

H	0.468080	4.277185	-1.220973

H	3.252668	6.532439	-1.170872

C	3.986201	3.126384	-1.914488

C	3.148702	3.414629	-3.163641

O	1.917520	2.674873	-3.155692

H	2.142302	1.727815	-3.217231

H	2.872225	4.469705	-3.226713

H	3.747349	3.154979	-4.047656

O	5.352703	2.903393	-2.332428

H	3.590602	2.205291	-1.459526

H	4.609152	5.074693	-1.250252

H	4.242272	2.737824	0.690849

H	6.718603	4.189041	-0.356668

H	7.148440	2.213904	-1.731187

C	5.003156	-0.840375	-2.128846

O	5.498979	-2.124338	-2.558147

H	4.173606	-0.977758	-1.417317

C	4.464167	-0.120966	-3.364663

O	3.031648	-0.061316	-3.397950

H	2.688318	-0.974838	-3.358476

H	4.811647	0.914310	-3.376799

H	4.855161	-0.624619	-4.260329

H	6.967339	0.101512	-2.078528

H	5.682440	-1.074050	0.429436

H	7.856514	-2.203918	-1.386718

H	6.255784	-3.896659	-1.986229

H	4.936094	-5.271759	-1.296653

C	2.985388	-3.970823	-3.057562

H	3.996255	-3.650114	-3.310201

H	2.670581	-4.734835	-3.780685

O	2.161809	-2.799001	-3.179272

H	1.226211	-3.062149	-3.124514

H	2.139767	-4.046163	-1.094348

H	1.837932	-7.594585	-0.861154

H	-0.215050	-7.428851	-0.788669

H	-1.236310	-4.535200	-0.796372

C	-0.519164	-5.312054	-2.662022

O	-0.659068	-3.965107	-3.132888

H	-1.599845	-3.715246	-3.066337

H	0.551076	-5.520607	-2.642569

H	-0.992189	-6.022727	-3.355473

H	-4.110137	-6.770861	-0.629922

H	-2.370957	-8.094326	0.367117

H	-1.118073	-5.515178	1.423148

H	-0.813372	-7.458804	2.674935

N	4.778139	0.303263	2.905987

H	4.469530	1.169281	3.385656

H	4.400686	0.376150	1.940564

C	4.284448	-0.952572	3.560635

C	2.782367	-1.147763	3.372151

H	2.509074	-2.002965	3.993607

H	2.247994	-0.278872	3.779185

C	2.307779	-1.431336	1.925316

C	2.048528	-0.142100	1.187731

O	2.983441	0.582619	0.797813

N	0.761553	0.218706	1.030839

C	0.370421	1.539270	0.547372

C	-0.909335	1.471391	-0.272322

H	-0.745821	0.830414	-1.150967

N	-2.462092	-0.933554	-0.146502

O	-1.654821	-1.224556	-0.955677

C	0.248817	2.516762	1.740451

O	-0.842044	2.967124	2.106863

N	1.414113	2.795925	2.340966

H	2.280894	2.408250	1.980621

H	3.068751	-1.992975	1.371322

H	1.402035	-2.044497	1.959691

H	4.485783	-0.841421	4.634298

C	5.140295	-2.154694	3.076969

O	6.290989	-1.893120	2.632289

O	4.622760	-3.288166	3.209302

H	5.799887	0.276589	2.864071

H	-1.183964	2.470792	-0.621050

H	0.797549	3.174417	4.315430

H	0.043834	-0.350952	1.462078

C	1.518147	3.549177	3.578731

H	1.285696	4.612141	3.415920

C	2.932880	3.470638	4.179528

O	3.845808	2.935598	3.461231

O	3.075581	3.957398	5.314305

S	-2.337889	0.778685	0.616278

H	1.173394	1.895974	-0.113088
